# Auxin signaling is a common factor underlying natural variation in tomato shade avoidance

**DOI:** 10.1101/031088

**Authors:** Susan M. Bush, Leonela G. Carriedo, Daniel Fulop, Yasunori Ichihashi, Mike F. Covington, Ravi Kumar, Aashish Ranjan, Daniel Chitwood, Lauren Headland, Daniele L. Filiault, José M. Jiménez-Goméz, Neelima R. Sinha, Julin N. Maloof

## Abstract

Light is an essential resource for photosynthesis. Limitation of light by shade from plant neighbors can induce a light competition program known as the shade avoidance response (SAR), thereby altering plant growth and development for the sake of survival. Natural genetic variation in SAR is found in plants adapted to distinct environments, including domesticated tomato *Solanum lycopersicum* and its wild relative *Solanum pennellii.* QTL mapping was used to examine variation of the SAR between these two species. We found organ specific responses in the elongation of the stem and petiole, including developmental acceleration of growth. Through RNAseq analysis we identified a number of ILs with reduced expression of auxin-related genes in shade treatment. These same ILs display a shade tolerant phenotype in stem growth and overall height. We also identified ILs with altered SAR expression of cell wall expansion genes, although these genotypes had no accompanying alteration in phenotype. Examination of weighted gene co-expression Connectivity networks in sun- and shade-treated plants revealed Connectivity changes in auxin and light signaling genes; this result was supported by the Identification of motifs within the promoters of a subset of shade-responsive genes that were enriched in light signaling, developmental pathways, and auxin responsive transcriptional domains. The Identification ofboth systemic and organ-specific shade tolerance in the ILs, as well as associated changes in the transcriptome, has the potential to inform future studies for breeding plants able to be grown closely (while neighbor-shaded), yet still maintaining high yield.

**Summary:** Growth plasticity in response to shade involves expression of specific auxin signaling and cell wall expansion genes, and shade avoidance QTL affect both stem elongation and developmental rate.

## Introduction

Plants depend on light for photosynthesis. Thus, for plants growing in densely populated stands, responding appropriately to nearby competitors is imperative for fitness (Dudley and Schmitt, 1996; Schmitt, 1997; Schmitt et al., 2003). Neighbor proximity is sensed via changes in the ambient light spectra. Chlorophyll preferentially absorbs red light (R), therefore light transmitted through or reflected from leaves becomes enriched with far-red light (FR); consequently, an alteration in the ratio of R and FR light is a signal of neighbor proximity. The spectral composition of shade (low R:FR light) induces a suite of transcriptional and developmental changes, known as the shade avoidance response (SAR), that promote plant growth through the elongation of internodes and petioles in an effort to gain better access to light (Casal, 2012).

Plants sense the environmental R:FR signal via the phytochrome family of photoreceptors (Mathews, 2010). Phytochromes are uniquely suited to the task of neighbor detection due to their capacity to photoconvert between inactive and active forms with absorption maxima in R (~660 nm) and FR (~730 nm) wavelengths. Hence, the pool of phytochrome proteins is in a dynamic equilibrium reflecting the relative proportions of R and FR in ambient light (Mancinelli, 1994). Activated phytochrome negatively regulates growth in high R:FR, ‘sun’, through its translocation from the cytosol to the nucleus. There, it interacts with and mediates degradation of the growth promoting PHYTOCHROME INTERACTING FACTORS (PIFs) (Nagy and Schafer, 2002; Khanna et al., 2004; Park et al., 2004). This family of bHLH transcription factors act as the primary hub for the SAR signaling cascade to promote elongation(Leivar and Quail, 2011). Because shade alters the spectral composition of R and FR, low R:FR, or ‘shade’, decreases the active pool of phytochrome, thereby allowing for the accumulation of PIFs and activation of their downstream targets.

Although SAR is an adaptive phenomenon varying within and between species (Morgan and Smith, 1979; Gilbert et al., 2001; Filiault and Maloof, 2012), the induction of a prolonged SAR can come at the cost of overall reproductive fitness (Dudley and Schmitt, 1996; Schmitt, 1997; Schmitt et al., 2003). Long-term exposure to low R:FR can result in a decrease in biomass, leaf area, and yield, and an acceleration of flowering (Donald Keiller, 1989; Devlin et al., 1996; Devlin et al., 1999; Cerdan and Chory, 2003). The domestication process of many crop species has resulted in a reduction of the SAR due to artificial selection for increased biomass or yield under high density planting. However, the SAR is not completely eliminated, which can be problematic for biomass and yield production if resources are shifted to stem elongation under crowded conditions (Casal and Kendrick, 1993; Skinner and Simmons, 1993; Page et al., 2010; Casal, 2012; Deng etal., 2012).

Because SAR can have negative effects on yield, the SAR in crops has mostly been studied at the physiological level, with a limited view of the genetic and transcriptional regulation of SAR. Our best genetic and molecular understanding of SAR is derived from studies of Arabidopsis, which provide a broad framework to better understand SAR in other species, including crops. Knowledge of the control of SAR in crops may be useful in the development of shade tolerant cultivars designed for maintenance crop yield to maximize arable land. This study focuses on the physiological and molecular mechanisms of SAR in tomato, which is among the top ten commodities globally, with a gross worth of US$84 million, and is closely related to many agronomic relatives in the Solanaceae family, including potatoes and peppers (FAOSTAT).

Recently, Cagnola and colleagues (Cagnola et al., 2012) characterized tomato SAR at the transcriptional level. When comparing the transcriptomes of internodes and petioles, they found both shared and organ-specific responses to shade, with the greater response occurring in the internodes. They observed that low R:FR reduced the expression of genes involved flavonoid synthesis and isoprenoid metabolism. These transcriptional changes were in concordance with their observed physiological response, suggesting that low R:FR decreases photosynthetic capacity of internodes, which can be seen as a means to reduce the energetic cost of SAR in the stem (Cagnola et al., 2012).

This important work serves as a foundation for our exploration of the differential effects of low R:FR treatment on the growth responses in different species of tomato. Wild tomato species exhibit strong shade avoidance, while SAR in domestic tomatoes is attenuated and variable (SFig. 1). We have begun to investigate the genetic bases of these species differences by studying *Solanum pennellii,* a species that is much more shade responsive than domesticated tomato. We exploited this interspecific difference by using a genetic population developed for the purpose of quantitative trait loci (QTL) Identification and derived from a cross between the cultivated tomato *Solanum lycopersicum* cultivar M82 (LA3475) and the wild, desert-adapted relative, *S. pennellii* (LA0716) (Eshed and Zamir, 1995). Within the introgression population, the *S. pennellii* genome is represented in 76 individual introgression lines (ILs), each bearing a small but overlapping segment of the wild species genome. Because the remainder of the genome in each IL is identical to that of M82, any phenotypic differences found between the IL and M82 can be attributed to the genes contained in the *S. pennellii* introgression in that line. Our recent high-resolution genotyping of these lines has precisely defined the genes introgressed into each line, allowing Identification of candidate genes for mapped traits (Chitwood et al., 2013).

This particular IL population has been useful in understanding many biological phenomena, including tomato fruit, leaf, and root development, dissecting the genetic basis for biotic and abiotic stress tolerance, brix content, metabolic profiling and, as we have found, light sensitivity (Eshed and Zamir, 1995; Overy et al., 2005; Frary et al., 2010; Chitwood et al., 2012; Chitwood et al., 2013; Hiroki Ikeda, 2013; Ron et al., 2013; Sharlach et al., 2013; Chitwood et al., 2014). To expand our understanding of the transcriptional control of the SAR in tomato, we performed RNA sequencing of juvenile ILs and IL parents under simulated sun and short-term shade exposure. For our phenotypic QTL study, we measured growth of five-week-old plants grown under simulated sun and shade conditions.

Using the IL population, we have taken a QTL mapping approach to study the phenotypic and associated gene expression responses to shade in tomato. Taking a principal component approach to morphological traits, we identified organ-specific elongation and developmental acceleration. Furthermore, our phenotypic analyses revealed a number of ILs with attenuated shade responses (shade tolerance), as well as shade sensitive ILs, in terms of growth and developmental acceleration. We also found a range of shade responsive gene expression patterns, some of which are consistent with Arabidopsis studies and the first tomato transcriptome findings, and we also identify novel tomato shade response genes.

## Results

The wild species *S. pennellii* shows a strong response to shade (Fig. 1A, SFig. 1), as might be expected from a plant adapted to an open, desert habitat. By contrast, the domesticated tomato *S. lycopersicum* cv. M82 has been bred for production in crowded field conditions, resulting in attenuated response to shade (Fig. 1B). Because the IL population is comprised of lines each carrying a small region of the *S. pennellii* genome in an M82 genetic background, they provide an excellent resource to search for the genetic underpinnings of shade response in tomato.

**Figure 1.**
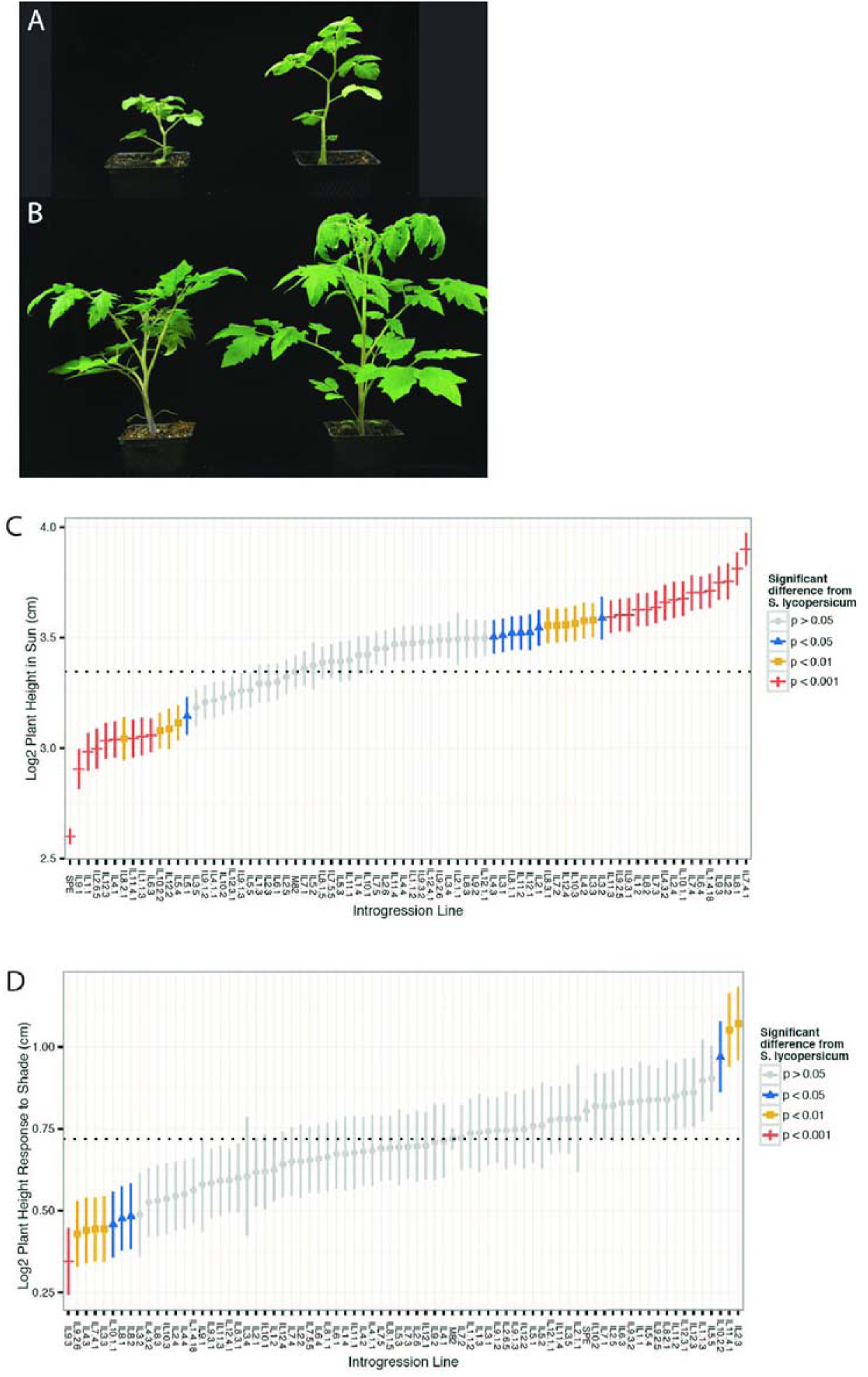
Domesticated and wild tomato species display variationariation in their growth responses to shade. (A) *S. pennellii* in high R:FR (sun, left) and low R:FR (shade, right). (B) *S. lycopersicum* cv. M82 in high R:FR (sun, left) and low R:FR (shade, right). (C) and (D) Log2-transformed height and Standard error of plants grown in sun (C) and change in height in response to shade (D), in the M82 × *S. pennellii* introgression population. Introgression lines are labeled on the x-axis.

### Tomato introgression lines show variation in many physical traits in response to shade

To dissect the genetic basis for variation in the tomato phenotypic response to shade, we grew the IL population under simulated sun (high R:FR = 1.5) and simulated shade conditions (low R:FR = 0.5). As previously stated, classic SAR includes the elongation of internodes and petioles. In order to identify genomic regions associated with variation in SAR, we measured a suite of traits known to be affected by low R:FR, including total height, internode length, petiole length, internode number, and flower number. Internode lengths were measured at two time points; comparison of the four- and five-week internode measurements permitted calculation of a growth rate for each IL in sun and shade. Across the ILs, we identified a broad range of growth variation including growth and developmental responses both greater and smaller than M82 (Fig. 1C-D).

### Principal component analysis identifies overall size and organ age as primary sources of variation in plant organ size

Analysis of the elongation measurements revealed strong correlations between many traits. For example, a plant with a long first internode was likely to have a long third internode (correlation of 0.39, *p*-value < 0.001; SFig. 2). To reduce the complexity of the data and to identify the major sources of variation in our morphological measurements, we utilized principal component analysis (PCA) (Wold et al., 1987). PCA transforms the raw, highly correlated data into orthogonal variables representing the primary sources of variation in the data. The principal components (PCs) can be treated similarly to raw trait measurements for downstream analyses. For example, we can identify genotypes with mean PC values significantly above or below the M82 mean, thereby identifying chromosomal regions that affect the trait or PC of interest. Internode measurements (epicotyl through internode 3), petiole measurements (leaf 1 through 4), and growth rate measurements (epicotyl through internode 3) were each independently subjected to PC analysis.

For both internode and petiole measurements, PC1 explains > 50% of the variation in the data (SFig. 3A). As shown in Fig. 2A-B, PC1 has similarly weighted loading for each internode or petiole, representing overall plant height (for internode measurements; intPC1) or overall petiole length (for petioles; petPC1) of the plant’s organs. In general, plants grown under shade have higher values of intPC1 and petPC1 because shade plants tend to be taller and have longer petioles than plants grown in sun (*t*-test *p*-value < 0.001; Fig. 2C). However, we found that a number of ILs had a differential organ response, as we highlight in Fig. 3. Fig. 3A presents the log_2_-transformed ratio of petPCl values (shade/sun). Large ratios indicate shade responsive genotypes (e.g. IL10.2), whereas low ratios indicate shade tolerance (e.g. IL9.1; see also SFig. 4).

**Figure 2.**
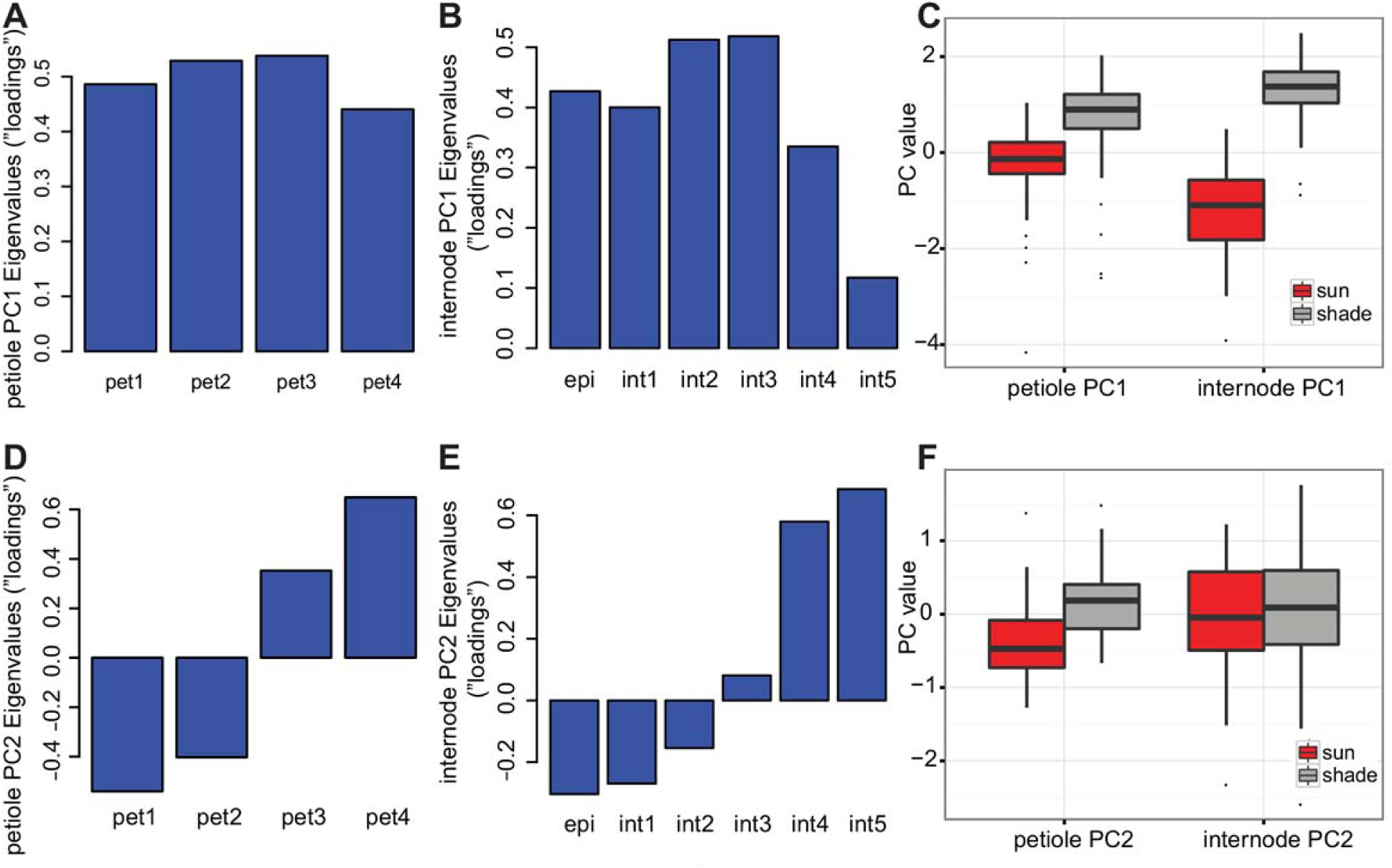
Principal component analysis of petiole and internode measurements. (A) and (B) PC1 represents 65% of variation in petioie length measurements (A) and 54% of variation in internode measurements (B), representing overall size of the organs. (C) Values for PC1 differentiate between plants grown in shade or sun; *t*-test *p*-values < 0.001. (D) and (E) PC2 represents 24% of variation in petiole length measurements (D) and 24% of variation in internodes (E), distinguishing between growth in older or younger organs. (F) Values for internode PC2 do not reliably distinguish between plants grown in shade or sun, although petiole PC2 does.

**Figure 3.**
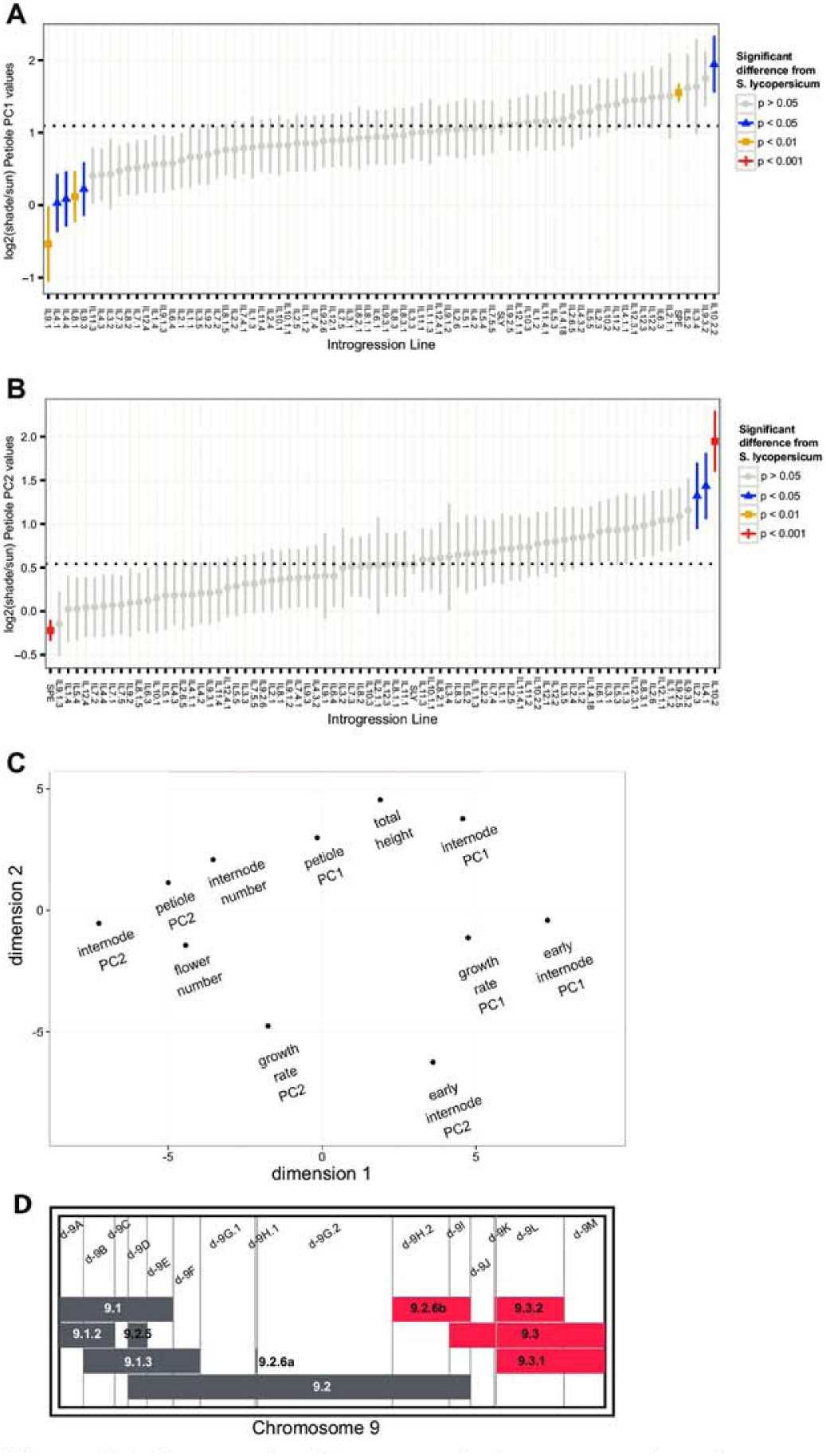
Introgression lines reveal chromosomal regions associated with shade tolerance and shade-responsive growth. (A) and (B) Means and standard errors of the relative shade response in petiole PC1 (A) and PC2 (B) values for each introgression line (log2-transformed ratios of shade/sun). *P*-values represent the significance compared to M82. (C) Multidimensional scaling plot of phenotypic traits and PCs presents the relationships between growth traits and organ age traits. (D) The chromosome 9 map of M82 × *S. pennellii* tomato introgression lines, with horizontal boxes representing regions of the S. pennellii genome in the indicated ILs <e.g. IL9.1). Vertical lines delineate bins as the unique and shared regions along the chromosome, defined by overlapping introgression regions (e.g. d-9A). Over-lapping ILs of interest are indicated in red.

PC2 explains approximately one quarter of the variation seen in the internode and the petiole data (SFig. 3A). The genotypes with high PC2 values are those that have relatively more growth in the younger tissues as compared to the older tissues, while a low PC2 suggests greater growth in the older organs such as the epicotyl or internode 1 (Fig. 2D-2E; SFig. 4C-D). In contrast to PC1, PC2 values do not vary greatly with light treatment (Fig. 2F), particularly in internodes; although *t*-test *p*-values are less than 0.001 for both intPC2 and petPC2, the dataset used for this analysis was sufficiently large (*n* = 1680) that there may not be biological significance underlying this statistical significance. PC2 values are closely related to the number of internodes present at the time of measurement, such that a high PC2 value associates with increased organ numbers (Spearman correlation, *rho* > 0.5, *p* < 0.001; SFig. 4I). Therefore, intPC2 can be thought of as representing the developmental rate of the plant, where a high value of intPC2 indicates the plant has progressed further in development than average. Similarly, petPC2 explains elongation of the petioles of younger versus older leaves. Fig. 3B shows the log_2_-transformed values of petiole PC2 (shade/sun). In this way, we can visualize those genotypes that have a significantly larger or smaller change in their development in shade, in comparison to the state of the plant grown in sun. In summary, PC2 allows the Identification of genotypes that alter the rate of their development in shade, with a large PC2 ratio indicating growth of more organs than M82 in shade (*e.g.* IL10.2, Fig. 3B).

Since SAR is known to accelerate growth in other species, we also asked how SAR impacts internode growth rate among the ILs. Plants were measured at 4 and 5 weeks of growth. The week 4 (“early”) measurements included epicotyl and internodes 1 to 3 (SFig. 3B-C); the log_2_-transformed ratio of shade/sun for early PCs 1 and 2 in the ILs are presented in SFig. 4E-F. Early PCs reveal those ILs that are shade tolerant at 4 weeks, a list that only partly overlaps with the shade tolerant ILs at week 5 (*e.g.* IL7.4.1, IL1.4.18). In order to determine the growth rate between weeks 4 and 5, week 4 measurements were subtracted from week 5 values for epicotyl and internodes 1 to 3. As for other traits, growth rate values were subjected to PC analysis, and the log_2_-transformed ratio of shade/sun for growth rate PC1 and PC2 are shown in SFig. 4G-H (see also SFig. 3D-E).

We then wanted to determine the relationship between the different PC traits classified as growth (PC1) or organ age (PC2) under shade. Specifically we were curious if the PCs would cluster by organ type or by trait type. A multidimensional scaling (MDS) approach showed that traits of overall size cluster near one another (total height, intPC1, petPC1) (Fig. 3C). The traits describing the age of developing organs or stage in development, including intPC2, petPC2, internode number, and flower number, also group together. Early and growth rate traits are clustered relatively apart from the terminal traits. Considering Fig. 3A-B and S4 Fig., the distinction of PC1 and PC2 traits can again be seen, where genotypes that are highly shade responsive in their overall growth do not necessarily increase their rate of development in shade (*e.g.* IL 11.4.1, IL8.2.1; S4 Fig. C-D). This Separation of PCs 1 and 2 can be seen across organs (intPC vs. petPC) and is maintained in both week four (early intPC) and week five (intPC) phenotypes, indicating separable genetic control for growth and developmental acceleration.

### Introgression lines display both systemic and organ specific responses to shade

The internode and petiole PC analyses allowed us to detect differences in growth (described by PC1) and development (as described by PC2), therefore allowing the Identification of a number of shade-sensitive and shade-tolerant ILs and those that are developmentally altered by shade (Fig. 3, SFig. 4). We noted that many ILs respond strongly to shade in internodes or in petioles but not necessarily both. The observation that the leaf and stem respond independently to shade suggests that SAR is regulated by many different loci that may control both systemic and organ specific responses to shade. The PC data was examined in order to determine whether specific ILs, therefore genomic regions or specific genetic loci, could be associated with the traits of interest.

An example of the differential growth and developmental response under shade described above can be highlighted by comparing the shade responses of two overlapping ILs with opposing phenotypes spanning a region on chromosome 9: IL9.3 and the overlapping IL9.3.2, bearing 400 fewer *S. pennellii* genes (Fig. 3D; overlapping ILs are indicated in red). IL9.3 is shade tolerant in both its internodes and petioles (Fig. 3A, SFig. 4A, B), and despite this line being shorter than M82 in shade, it does not show a developmental rate difference in shade (Fig. 3B, SFig. 4D). By contrast, IL9.3.2 has a greater number of internodes and more growth occurring within the younger tissues, represented by high PC2 values, suggesting this line accelerates its growth under shade (Fig. 3B; SFig. 4A, D). The fact that the IL9.3.2 introgressed region is contained entirely within IL9.3 and displays growth acceleration in shade suggests that the additional genes in IL9.3 inhibit growth. A subset of the *S. pennellii* genes in IL9.3 may act as negative regulators of developmental acceleration under shaded conditions, while the M82 alleles of those genes present in IL9.3.2 do not. The relationship between the phenotypes of IL9.3 and IL9.3.2 can also be seen in Fig. 4, a heatmap presenting the shade responses of each phenotype across the ILs.

**Figure 4.**
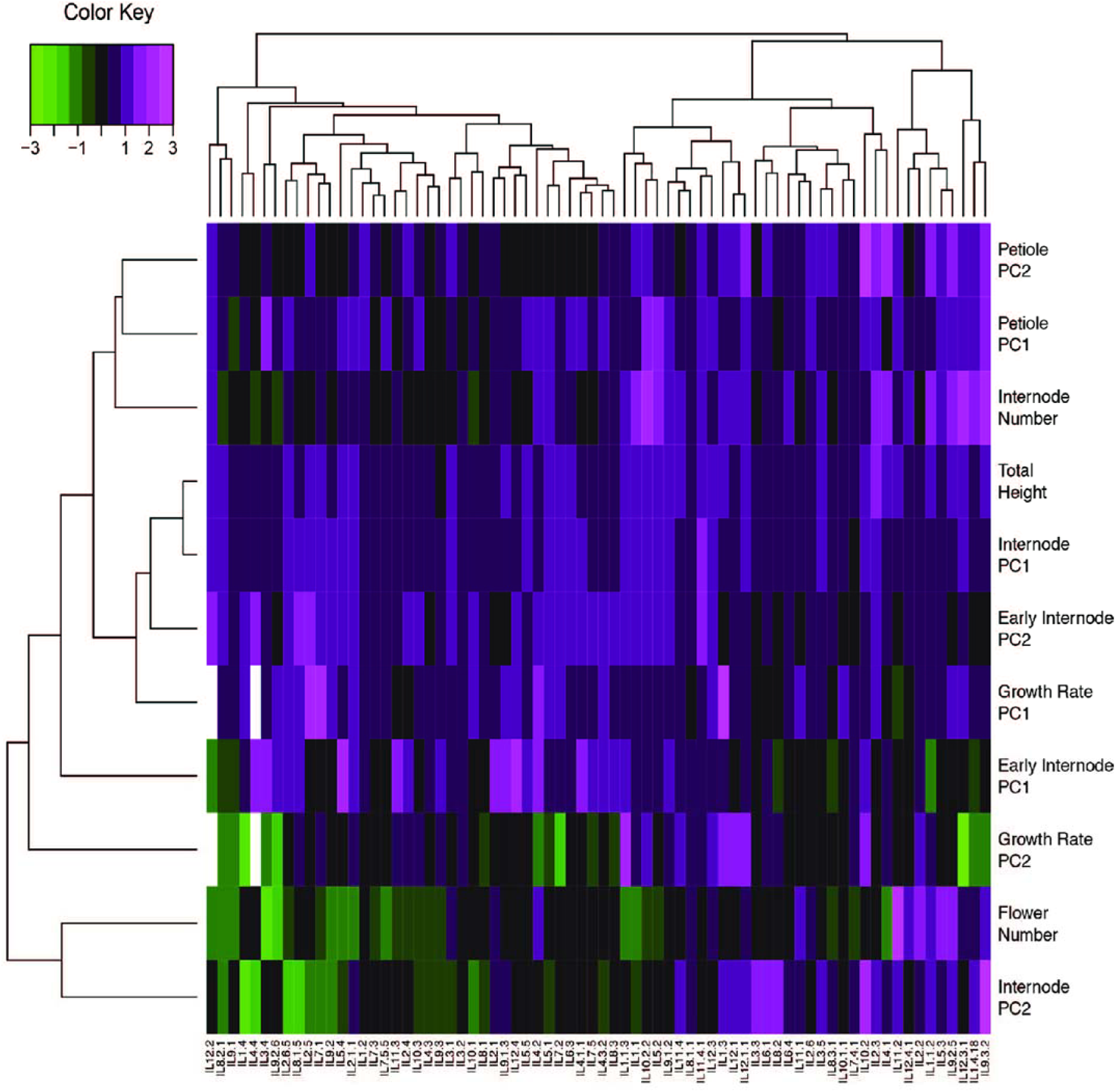
A heatmap displays the relative shade responses in each phenotype across the introgression lines, clustered by Euclidean distances. Relative shade response is calculated as the log2-transformed ratios of shade/sun values. Data were scaled within each phenotype prior to clustering. Green coloring represents traits where shade expression values are less than those in sun (values < 0); magenta coloring represents values > 0 where shade values are greater than those in sun. White values indicate missing data.

In the aforementioned example, we highlight a benefit of knowing the boundaries of the introgression lines in that it allows us to precisely define the “bins”, or the overlapping segments of each contiguous IL per chromosome. We used this bin mapping approach to ask whether a particular segment of an IL, or chromosome, contributes to a particular phenotype. We asked whether the loci negatively regulating developmental rate in IL9.3 also controlled the SAR observed in the petioles. The data revealed that none of the IL9.3 neighbors showed a differential shade avoidance response in petioles, thus allowing us to hypothesize that the location of these antagonizing regulators must be in bin d-9J, the only bin found uniquely in IL9.3 (Fig. 3D).

Bin mapping allowed us to dissect lines with organ specific or systemic responses to shade. Compared to their neighboring chromosomal ILs, the ILs 7.4.1 and 3.2 are both significantly taller than M82 under sun conditions, yet are also significantly less shade responsive than M82 (Fig. 1C, SFig. 4C). Together, these results indicate that inherent growth characteristics (tall under normal circumstances) may limit how much additional growth is possible in response to shade, or that these lines display a constitutive shade avoidance phenotype. In contrast, IL11.4.1 is significantly shorter than M82 in sun, and it has a robust shade response in overall height; however, its petiole elongation and growth rate do not display SAR (SFig. 4B-C versus Fig. 3A-B). An additional example of organ specific SAR is seen in IL2.3, which possesses a significant shade response for internode length and height but not for petioles (Fig. 3A-B, SFig. 4B-C). These findings suggest that some of the mechanisms controlling elongation under shade are organ specific.

This manual IL-by-IL bin mapping approach is classically used in IL studies to determine bin-to-phenotype causality. However, it is not practical when attempting to understand the complexity of the SAR demonstrated among a large population and for several phenotypes. Therefore, we used a computational method, elastic net regularized regression (Zou and Hastie, 2005) to examine the contributions of each bin within an introgression to the shade responsive phenotypes, and to simultaneously map the relative effect of each bin in the genome (Fig. 5; see also SFig. 5). This elastic net bin-mapping approach demonstrated that each phenotype in sun and shade is controlled by a dynamic set of QTLs (Fig. 5, SFig. 5). This approach not only validated our earlier findings that bin d-9J may contain a negative regulator of the SAR in petioles, it also expanded our view of which additional bins may be playing a role in regulating the SAR in tomato (Fig. 5, SFig. 5D, F).

**Figure 5.**
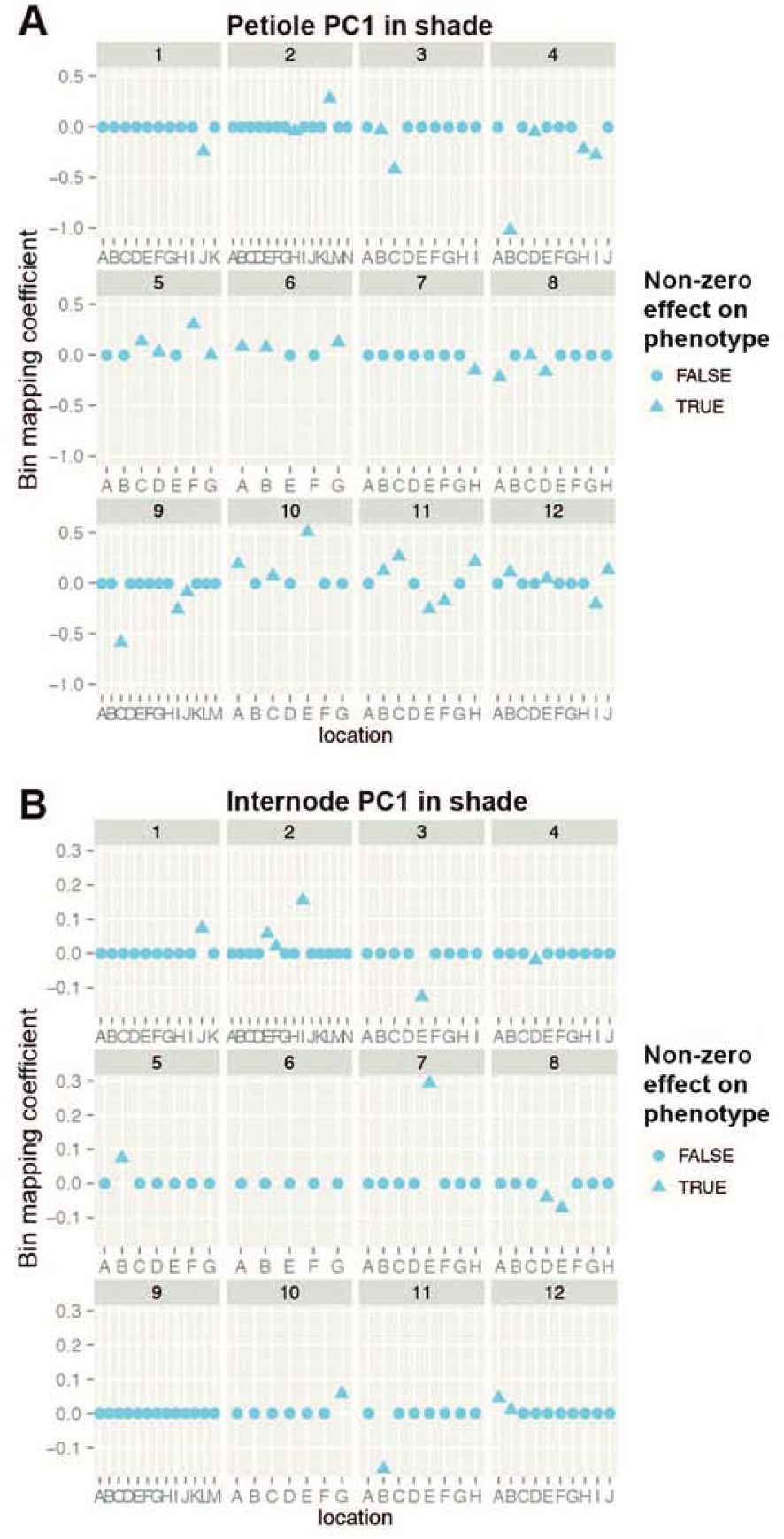
Shade-responsive traits are mapped to bins within the introgression lines. We determined the contribution of each bin to a given phenotype using elastic net regularized regression. A value equal to 0 indicates the bin does not contribute to the phenotype. The contributions of each bin to petiole PC1 in shade (A) or internode PC1 in shade (B) are shown, with circles representing no effect on phenotype and triangles representing a non-zero effect.

While our previous analyses allowed us to map or correlate traits to specific ILs, bin mapping allows us to focus on a smaller subset of genes that may be affecting each trait. For example, Fig. 3A indicates that IL4.1 has a weak shade response in petiole PC1; Fig. 5A demonstrates that that is primarily due to bin d-4B and not the other four bins present in IL4.1. The introgression bins d-9I and d-9J, present in IL9.3 but not in IL9.3.2 (Fig. 3D), can be seen to have a negative effect on petiole PC1, or growth in petioles, as suggested by our previous analysis; the bin mapping supports these results. Bin-mapping also confirmed the internode PC1 shade response we reported for IL2.3 (SFig. 4C); Fig. 5B indicates the response is due to contributions from three bins: d-2E, d-2F, and d-2I. This method can help narrow down the number of possible candidate genes underlying variation in SAR; the genes found within these chromosome 2 bins are listed in STable 1.

### Tomato expresses both known and novel genes in response to growth in shade

Our analysis of phenotypes in the IL population was undertaken to identify regions of the genome that are involved in shade-responsive phenotypes. To explore the underlying cause of the phenotypic variation seen in the SAR in the IL population, we sequenced the transcriptomes from the apical meristem of young plants as described by Kumar *et al.* (Kumar et al., 2012). Plants were grown in sun then shifted to growth in either sun (high R:FR) or shade (low R:FR) 28h prior to tissue collection.

Similar to the physical phenotypes described earlier, we measured gene expression in sun conditions as well as the change in expression in response to growth in shade. Using transcriptome sequence Information from each IL in sun and in shade, we performed differential gene expression analyses between genotypes and between light treatments using edgeR (Robinson et al., 2010). In this manner, we were able to identify a pool of shade-responsive genes in tomato, as well as genes up- or down-regulated in each IL in response to shade.

We looked at the top 8,000 genes differentially expressed under shade to determine whether previously reported shade responsive genes were also involved in the tomato SAR (STable 2). In table 1, we show the top 50 most differentially expressed genes across the IL population, irrespective of genotype, in response to 28h of simulated shade (Table 1). We found three tomato homologs of *ATHB2* (for *ARABIDOPSIS THALIANA HOMEOBOX PROTEIN 2*) and a number of *XYLOGLUCAN ENDO-TRANSGLYCOSYLASE/HYDROLASE* (*XTH*) genes, as well as genes involved in auxin and GA signaling (Table 1; (Carabelli et al., 1993; Sessa et al., 2005; Pierik et al., 2009; Kozuka et al., 2010). As shown in Table 1, *ATHB2* expression is increased in shade-treated plants, as is auxin-related gene expression and *XTH* expression. Expression of *GID1* (for *GIBBERELLIN INSENSITIVE DWARF*), the GA receptor, is lowered in shade-treated plants (Table 1; STable 2). We also identified several genes that had not previously been shown to be a part of phytochrome B-mediated shade avoidance. These novel genes include the tomato homologs of *FAR-RED INSENSITIVE 219/JASMONATE RESISTANT 1,* shown in Arabidopsis to be involved in *CONSTITUTIVE PHOTOMORPHOGENIC 1* (*COP1*) light-regulated signaling (Hsieh et al., 2000) and in phytochrome A low light responses (Robson et al., 2010). The metabolic gene *ATP CITRATE LYASE/SYNTHASE A 2* (*ACLA-2*) is increased in expression in shade-treated plants; other members of the citrate lyase pathway are also up-regulated by shade, but at a lower level. With its important role in synthesis of the citric acid cycle precursor acetyl-CoA, *ACLA-2* has been shown to be critical for growth in Arabidopsis (Fatland et al., 2002; Fatland et al., 2005).

**Table 1.** Top 50 genes identified as most significantly responsive to shade in tomato.

| ITAG | logFC in shade | logCPM | PValue | FDR | AGI | Gene name | Gene description |
| --- | --- | --- | --- | --- | --- | --- | --- |
| Solyc08g078300 | 0.84 | 4.14 | 7.29E-190 | 1.31E-185 | AT4G16780 | ATHB-2 | ATHB-2 (Homeobox-leucine zipper protein HAT4); DNA binding / transcription factor |
| Solyc06g060830 | 0.87 | 4.44 | 1.59E-186 | 1.43E-182 | AT4G16780 | ATHB-2 | ATHB-2 (Homeobox-leucine zipper protein HAT4); DNA binding / transcription factor |
| Solyc06g067910 | 1.16 | 4.73 | 3.00E-169 | 1.80E-165 | AT5G11420 | NA | unknown protein |
| Solyc06g008580 | 1.01 | 3.23 | 7.15E-113 | 3.21E-109 | AT5G43700 | ATAUX2-11 | Auxin inducible transcription factor |
| Solyc01g110660 | 0.87 | 3.15 | 9.00E-111 | 3.24E-107 | NA | NA | NA |
| Solyc08g066740 | 0.89 | 5.55 | 4.83E-108 | 1.45E-104 | NA | NA | NA |
| Solyc07g017600 | 0.48 | 5.14 | 5.30E-105 | 1.36E-101 | AT4G33220 | ATPME44 | Pectinesterase family protein |
| Solyc10g083760 | -0.25 | 6.97 | 1.27E-104 | 2.85E-101 | AT3G10050 | OMR1 | Enzyme in the biosynthetic pathway of isoleucine |
| Solyc10g039290 | 0.60 | 7.75 | 3.43E-103 | 6.85E-100 | NA | NA | NA |
| Solyc10g076820 | -0.76 | 3.05 | 9.94E-100 | 1.79E-96 | AT2G38090 | NA | Myb family transcription factor |
| Solyc05g013440 | -0.47 | 7.28 | 1.06E-95 | 1.74E-92 | AT2G42490 | NA | Copper amine oxidase, putative |
| Solyc07g049380 | 1.32 | 1.28 | 2.59E-93 | 3.88E-90 | NA | NA | NA |
| Solyc03g096840 | -0.41 | 7.34 | 6.75E-90 | 9.33E-87 | NA | NA | NA |
| Solyc06g075440 | -0.88 | 1.89 | 1.48E-89 | 1.90E-86 | NA | NA | NA |
| Solyc04g007690 | 0.88 | 3.89 | 2.46E-88 | 2.95E-85 | AT1G70940 | ATPIN3 | Regulator of auxin efflux, involved in differential growth, expressed in gravity-sensing tissues |
| Solyc06g008590 | 0.87 | 4.56 | 1.05E-87 | 1.18E-84 | AT3G04730 | IAA16 | Early auxin-induced (IAA16) |
| Solyc06g075510 | -0.40 | 5.30 | 1.69E-87 | 1.79E-84 | AT2G28550 | RAP2.7 | TOE1 (TARGET OF EAT1 1); DNA binding / transcription factor |
| Solyc02g092000 | 0.32 | 5.69 | 1.92E-86 | 1.92E-83 | AT3G49720 | NA | unknown protein; contains domain S-adenosyl-L-methionine-dependent methyltransferases |
| Solyc12g010800 | 0.68 | 3.75 | 5.58E-86 | 5.27E-83 | AT2G42380 | ATBZIP34 | bZIP transcription factor family protein |
| Solyc12g007230 | 0.37 | 5.98 | 1.20E-84 | 1.08E-81 | AT4G29080 | IAA27 | Phytochrome-associated protein 2 (PAP2) |
| Solyc11g010100 | 0.35 | 5.26 | 1.87E-83 | 1.60E-80 | AT5G65270 | AtRABA4a | Arabidopsis Rab GTPase homolog A4a); GTP binding |
| Solyc01g111350 | -0.44 | 7.58 | 1.09E-81 | 8.86E-79 | AT4G34950 | NA | Nodulin family protein |
| Solyc06g005610 | 0.51 | 2.99 | 1.18E-81 | 9.24E-79 | NA | NA | NA |
| Solyc01g102720 | 0.65 | 4.83 | 1.66E-81 | 1.24E-78 | AT5G49800 | NA | unknown protein with Lipid-binding START |
| Solyc06g008870 | -0.54 | 3.53 | 9.54E-80 | 6.85E-77 | AT3G63010 | ATGID1B | Gibberellin receptor (GID1). Interacts with DELLA proteins in vivo in the presence of GA4. |
| Solyc08g007270 | 0.62 | 3.24 | 1.30E-79 | 8.97E-77 | AT4G16780 | ATHB-2 | ATHB-2 (Homeobox-leucine zipper protein HAT4); DNA binding / transcription factor |
| Solyc09g065560 | -0.57 | 3.53 | 1.03E-78 | 6.85E-76 | AT3G51895 | AST12 | Encodes a sulfate transporter. |
| Solyc03g118750 | 0.67 | 6.04 | 2.69E-78 | 1.73E-75 | AT3G18000 | NMT1 | Arabidopsis thaliana N-methyltransferase-like protein |
| Solyc12g094380 | -0.63 | 5.22 | 2.68E-77 | 1.66E-74 | AT1G76020 | NA | unknown protein, contains Thioredoxin fold |
| Solyc12g011030 | 0.95 | 3.55 | 4.04E-77 | 2.42E-74 | AT4G14130 | XTH15 | Xyloglucan endotransglycosylase-related protein |
| Solyc01g095580 | -0.26 | 5.37 | 4.65E-77 | 2.69E-74 | AT2G46370 | FIN219 | An auxin-induced gene in GH3 family |
| Solyc09g083280 | 0.57 | 6.33 | 8.31E-76 | 4.67E-73 | AT5G43700 | ATAUX2-11 | Auxin inducible transcription factor |
| Solyc04g082270 | 0.78 | 3.12 | 1.30E-75 | 7.10E-73 | NA | NA | NA |
| Solyc03g123670 | 0.70 | 1.51 | 1.18E-74 | 6.23E-72 | NA | NA | NA |
| Solyc07g052320 | 0.37 | 4.62 | 7.50E-74 | 3.85E-71 | AT1G05170 | NA | Galactosyltransferase family protein |
| Solyc09g065100 | -0.59 | 3.74 | 1.29E-73 | 6.46E-71 | NA | NA | NA |
| Solyc08g021820 | 0.94 | 1.83 | 8.65E-73 | 4.20E-70 | NA | NA | NA |
| Solyc12g008940 | 0.17 | 8.21 | 3.46E-72 | 1.64E-69 | AT2G19480 | NAP1;2 | Nucleosome Assembly Protein1;2 DNA binding |
| Solyc07g049450 | 0.18 | 7.33 | 2.20E-71 | 1.01E-68 | AT1G04980 | ATPDI10 | Protein disulfide isomerase-like (PDIL) protein in the thioredoxin (TRX) superfamily. |
| Solyc03g025190 | -0.64 | 6.25 | 4.21E-70 | 1.89E-67 | AT4G25640 | ATDTX35 | MATE efflux family protein, similar to Multi antimicrobial extrusion protein MatE |
| Solyc03g095310 | 0.50 | 4.23 | 2.62E-68 | 1.15E-65 | AT5G24910 | CYP714A1 | Member of CYP714A |
| Solyc02g085020 | -0.66 | 7.17 | 3.30E-68 | 1.41E-65 | AT5G42800 | DFR | Dihydroflavonol reductase in anthocyanin synthesis |
| Solyc01g110630 | 1.05 | 0.75 | 8.53E-68 | 3.56E-65 | AT4G38840 | NA | Auxin-responsive protein, putative similar to auxin-induced SAUR-like |
| Solyc03g118040 | 0.23 | 9.26 | 1.57E-67 | 6.42E-65 | AT5G61790 | ATCNX1 | Calnexin 1 (CNX1) |
| Solyc01g058390 | -0.17 | 6.05 | 6.18E-67 | 2.47E-64 | AT3G06580 | GAL1 | Encodes a protein with galactose kinase activity. |
| Solyc01g087560 | 0.20 | 7.37 | 7.07E-67 | 2.76E-64 | AT5G13710 | CPH | SMT1 controls the level of cholesterol in plants |
| Solyc09g075450 | 0.23 | 7.07 | 2.25E-66 | 8.60E-64 | AT5G50950 | FUM2 | Fumarate hydratase, putative |
| Solyc04g079860 | 0.55 | 2.95 | 5.01E-66 | 1.88E-63 | AT3G50760 | GATL2 | Encodes a protein with putative galacturonosyltransferase activity. |
| Solyc12g094460 | 1.07 | 1.35 | 8.36E-66 | 3.06E-63 | AT1G76160 | sks5 | SKS5 (SKU5 Similar 5); copper ion binding / oxidoreductase; similar to SKS6 pectinesterase |
| Solyc04g009900 | -0.38 | 4.72 | 2.41E-65 | 8.66E-63 | AT1G08650 | ATPPCK1 | Phosphoenolpyruvate carboxylase kinase expressed in leaves and induced by light. |

### Shade-responsive gene expression across the tomato ILs is comparable to that found in Arabidopsis and tomato shade microarrays

To classify the types of genes regulated by shade, we tested for enrichment of gene ontology (GO) categories in the list of shade-regulated genes. The shade-enriched biological-process GO terms are given in Table 2. We used the gene ontologies to identify classes of genes contributing to the shade avoidance phenotypes. In a similar fashion, we compared the genes differentially expressed in our dataset with those identified in genetic pathways affected by shade. Using published gene sets from *Arabidopsis thaliana* (STable 4), we found tomato homologs of previously identified genes involved in flowering, cell wall modification, and light response pathways (Jiao et al., 2005; Mouhu et al., 2009; Koenig et al., 2013), as well as transcription factors (Davuluri et al., 2003) and genes regulated by hormones such as auxin, jasmonic acid, and gibberellin (Nemhauser et al., 2006). In concurrence with previous studies, one-third of the genes differentially expressed in our dataset have previously known to be light regulated (SFig. 7A). Genes previously shown to be shade responsive in Arabidopsis make up 8.5% of the genes differentially regulated by shade in tomato. Another third of the tomato shade-responsive genes were identified as Arabidopsis homologs that were not present in any of the lists we surveyed (SFig. 7A).

**Table 2.** Gene ontology biological process categories over- or under-represented in tomato shade-responsive genes.

| GO category | Biological Process GO term | Differential expression of GO category | p-value | Adjusted p-value |
| --- | --- | --- | --- | --- |
| GO:0042254 | ribosome biogenesis | up | 6.49E-73 | 1.69E-69 |
| GO:0006412 | translation | up | 1.08E-64 | 2.80E-61 |
| GO:0006886 | intracellular protein transport | up | 2.56E-31 | 6.66E-28 |
| GO:0016192 | vesicle-mediated transport | up | 9.98E-19 | 2.59E-15 |
| GO:0046686 | response to cadmium ion | up | 2.05E-18 | 5.32E-15 |
| GO:0015031 | protein transport | up | 8.88E-17 | 2.31E-13 |
| GO:0007264 | small GTPase mediated signal transduction | up | 3.83E-12 | 9.94E-09 |
| GO:0006184 | GTP catabolic process | up | 6.04E-12 | 1.57E-08 |
| GO:0006446 | regulation of translational initiation | up | 2.63E-11 | 6.82E-08 |
| GO:0018279 | protein N-linked glycosylation via asparagine | up | 1.37E-08 | 3.56E-05 |
| GO:0006913 | nucleocytoplasmic transport | up | 2.67E-08 | 6.90E-05 |
| GO:0032259 | methylation | up | 3.37E-08 | 8.73E-05 |
| GO:0006888 | ER to Golgi vesicle-mediated transport | up | 1.00E-07 | 0.000259511 |
| GO:0006413 | translational initiation | up | 4.94E-07 | 0.001274944 |
| GO:0006096 | glycolysis | up | 8.78E-07 | 0.002265101 |
| GO:0006094 | gluconeogenesis | up | 1.07E-06 | 0.002749404 |
| GO:0009955 | adaxial/abaxial pattern specification | up | 1.28E-06 | 0.003290272 |
| GO:0006457 | protein folding | up | 6.81E-06 | 0.017547984 |
| GO:0006461 | protein complex assembly | up | 9.42E-06 | 0.024281049 |
| GO:0006626 | protein targeting to mitochondrion | up | 1.47E-05 | 0.037801512 |
| GO:0000059 | protein import into nucleus, docking | up | 1.72E-05 | 0.044287923 |
| GO:0005983 | starch catabolic process | dn | 7.77E-11 | 2.01E-07 |
| GO:0009069 | serine family amino acid metabolic process | dn | 1.97E-08 | 5.09E-05 |
| GO:0006468 | protein phosphorylation | dn | 2.61E-08 | 6.75E-05 |
| GO:0006355 | regulation of transcription, DNA-dependent | dn | 3.61E-08 | 9.33E-05 |
| GO:0016310 | phosphorylation | dn | 1.23E-07 | 0.000318988 |
| GO:0016567 | protein ubiquitination | dn | 1.71E-07 | 0.00044298 |
| GO:0009773 | photosynthetic electron transport in photosystem I | dn | 2.97E-07 | 0.00076639 |

We wanted to further compare and contrast our tomato SAR gene expression results with that of previously reported Arabidopsis SAR gene expression findings. To that end, we utilized two well-known ATH1 microarray datasets in the Arabidopsis field released by Sessa and others (2005) and Tao and others (2008). We subset our tomato gene list to include only those genes with Arabidopsis homologs present on the microarray. The genes identified by Tao *et al.* (2008) and by Sessa *et al.* (2005) as being differentially expressed in shade in Arabidopsis representjust over 16% of the microarray-equivalent genes in our tomato shade data (SFig. 7B; enriched GO terms listed in STable 5). Not only does this point to inherent species level differences in the SAR, but also to the stark differences in the stage of development sampled in both these studies. The data published by Tao and colleagues was derived from 7-day-old Arabidopsis seedlings grown on plates and treated with 1h of shade, while the work done by Sessa and colleagues used 8-day-old seedlings treated with 1h or 4d shade. In contrast, our study addresses the early changes in gene expression due to shade exposure in a crop plant, using 19-day-old tomatoes grown on soil and treated with 28h-simulated shade. Additionally, our study utilized over 80 samples in each sun and shade treatment; compared to the Arabidopsis studies using up to 3 samples in each sun and shade, our study had much greater power to detect differential gene expression, which is likely reflected in the large number of genes identified.

### Gene expression patterns identify down-regulation of auxin-related genes in ILs with reduced SAR

In order to visualize the similarities and differences in shade-responsive gene expression in each IL, we clustered each genotype based on its expression of the ~ 8,000 genes differentially expressed in shade (SFig. 6). Fig. 6 presents a subset of shade-responsive genes, showing a heatmap of the ILs clustered based on the top 500 genes most differentially expressed in shade. These two heatmaps show distinct subsets of genes and ILs with related expression patterns, thus indicating trends in shade response. To ask whether changes in gene expression relate to differences in phenotype, we clustered the ILs based on their expression patterns in subsets of genes, including transcription factors, cell wall-associated genes, and auxin-responsive genes (SFig. 8). Looking at a heatmap of the subset of transcription factors, expression levels of the tomato homologs of *ATHB2* are increased in response to shade in nearly all ILs (SFig. 8A, highlighted box), as we expect from studies in Arabidopsis (Sessa et al., 2005; Tao et al., 2008) (genes listed in STable 4). Additionally, nine other transcription factors were universally up-regulated across the ILs in shade (SFig. 8A). Of these transcription factors, the Arabidopsis homolog of *ETHLYENE AND SALT INDUCIBLE 3* (*ESE3;* Solyc06g065820), was identified by Sessa *et al.* (2005) and Tao *et al.* (2008) as being up-regulated in shade; *ESE3* is known to be gibberellin and ethylene responsive. Expression of the other seven transcription factors, including *SUPPRESSOR OF OVEREXPRESSION OF CO1* (*SOC1*), *LOB DOMAIN-CONTAINING PROTEIN 1* (*LBD1*), and *FAMA,* have not previously been shown to be regulated by shade (Table 3).

**Figure 6.**
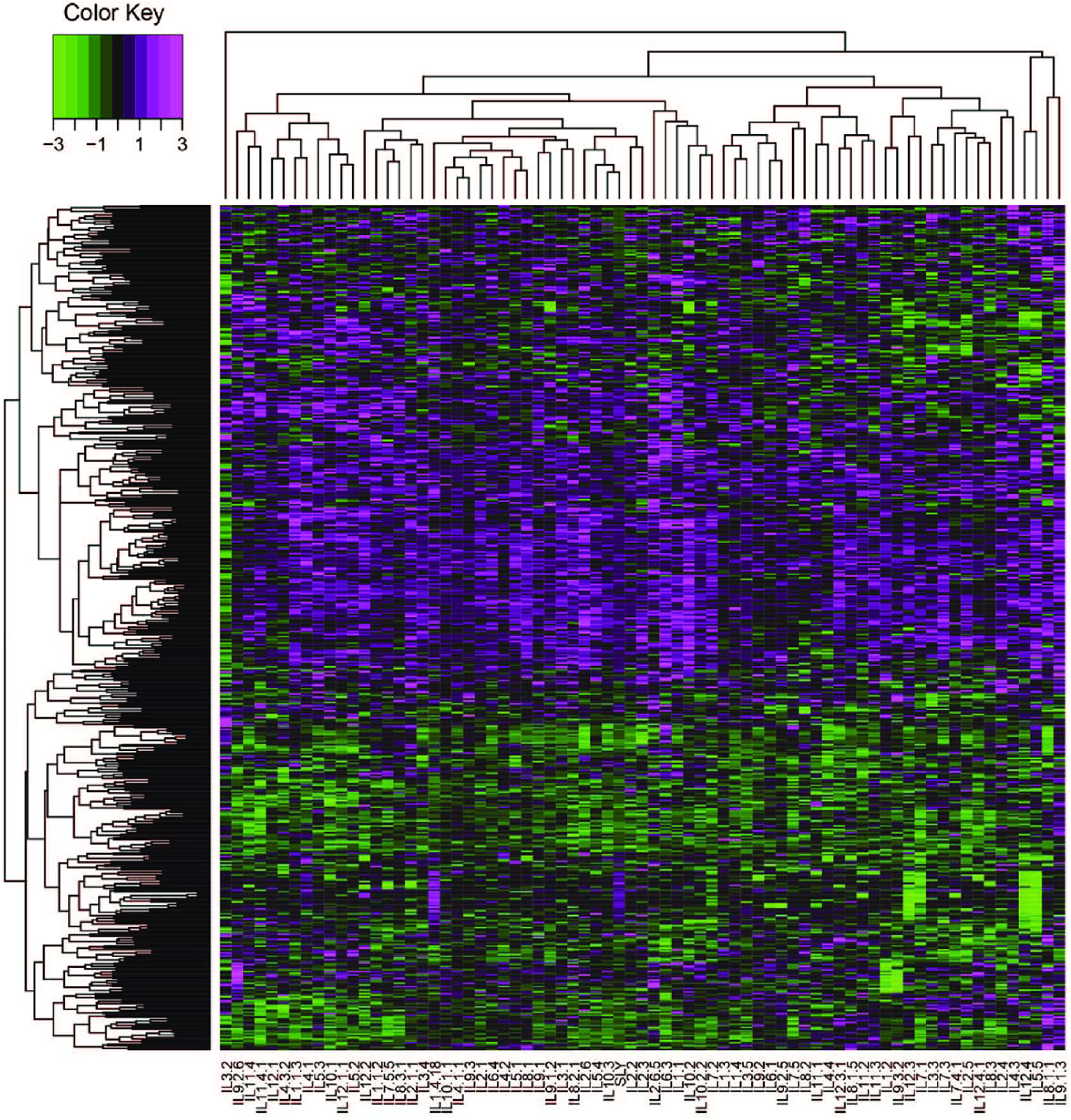
A heatmap displays the relative expression of the 500 most shade responsive genes across the M82 × *S. pennellii* tomato introgression lines, clustered by Euclidean distance. Relative expression is calculated as the log2-transformed ratios of shade/sun expression values. Data were scaled within each IL prior to ciustering. Green coloring represents genes where shade expression is less than that in sun (values < 0); magenta coloring represents genes > 0 where shade expression is greater than that in sun.

**Table 3.**
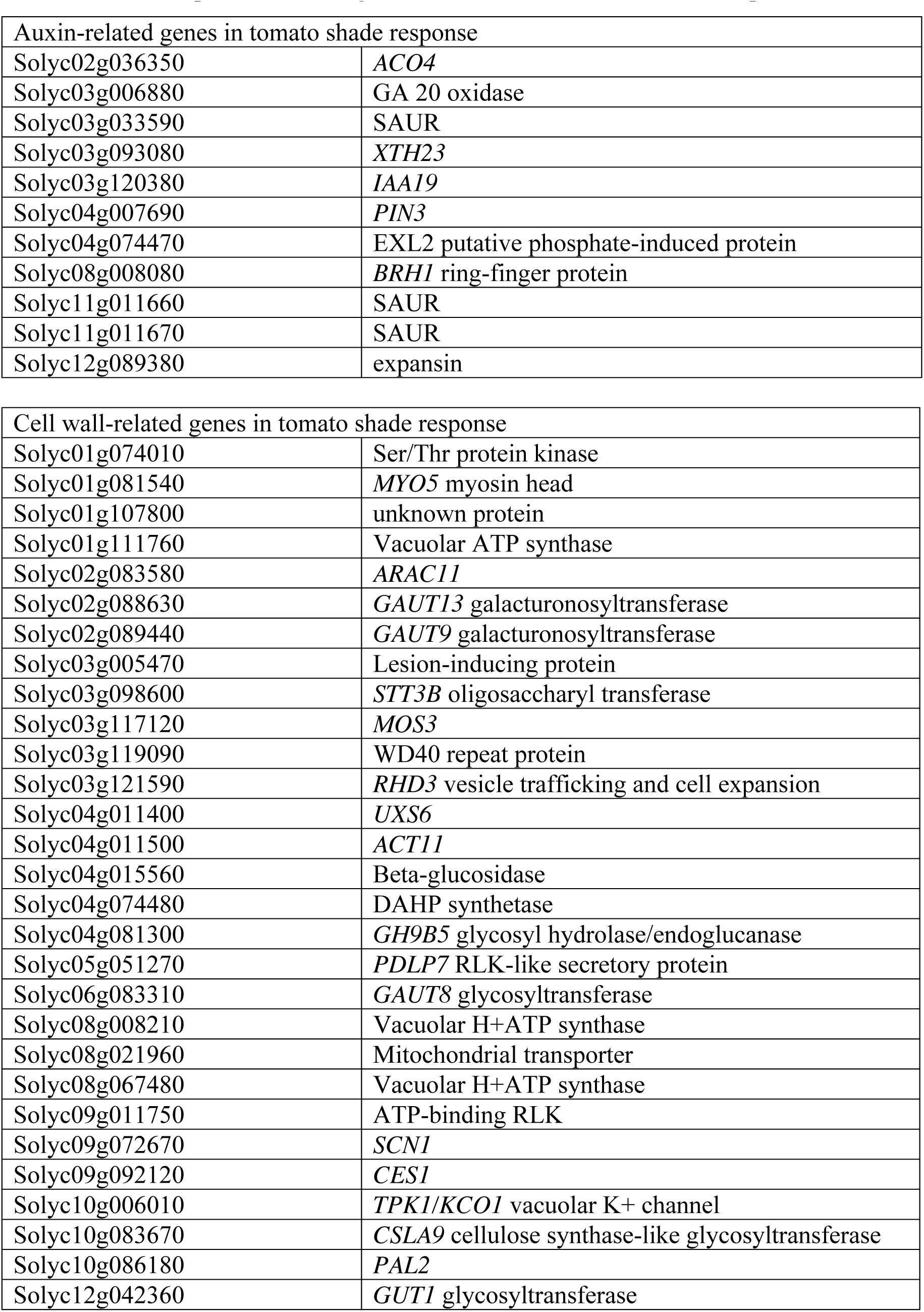
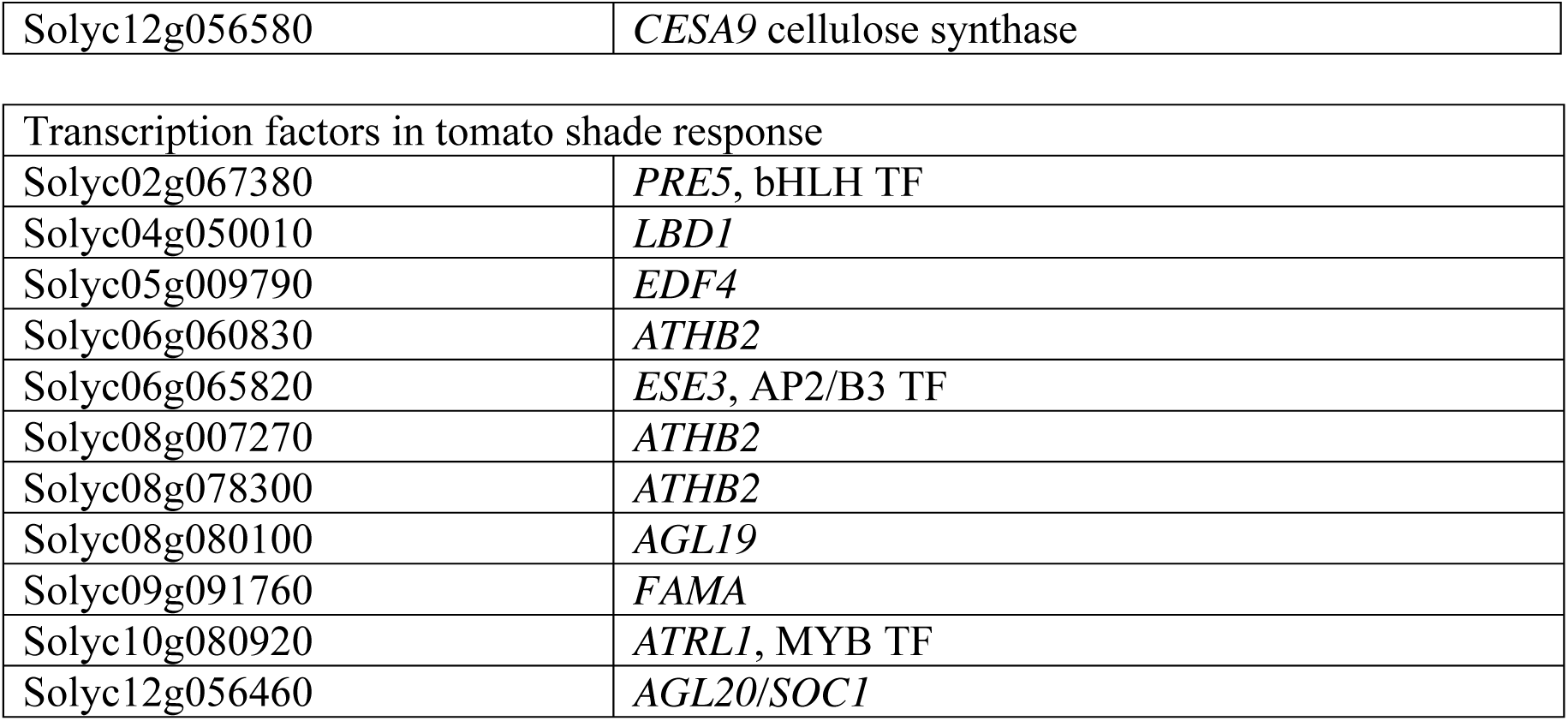
Shade-responsive tomato genes ofinterest identified based on expression in ILs

Shade-responsive expression of the cell wall-associated genes shows variation across the ILs. In particular, one group of ILs displays an atypical expression pattern with unchanged or reduced expression of approximately ten genes whereas most ILs show increased expression (SFig. 8B, highlighted box; Table 3); however, none of these introgression lines were identified as having a growth phenotype distinct from M82 in shade based on our measurements. In future experiments, it may be of interest to ask if the cell-wall composition in response to shade is different in this subset of ILs. The auxin-associated gene cluster in Fig. 7 (see also SFig. 8C) reveals a subset of ILs that have decreased expression of approximately ten auxin-responsive genes in shade (Table 3; STable 12); this set of ILs is included in those with a significantly reduced growth response in shade (Fig. 7, highlighted box; see also Fig. 1, SFig. 4B-C). In summary, changes in expression of cell wall-associated genes do not reflect the phenotypic differences we observed in the ILs, while differential expression of several auxin-related genes does, indeed, relate directly to the phenotypes measured in this study. This indicates that regulation of auxin signaling is a key convergence point for QTL Controlling shade avoidance in this mapping population.

**Figure 7.**
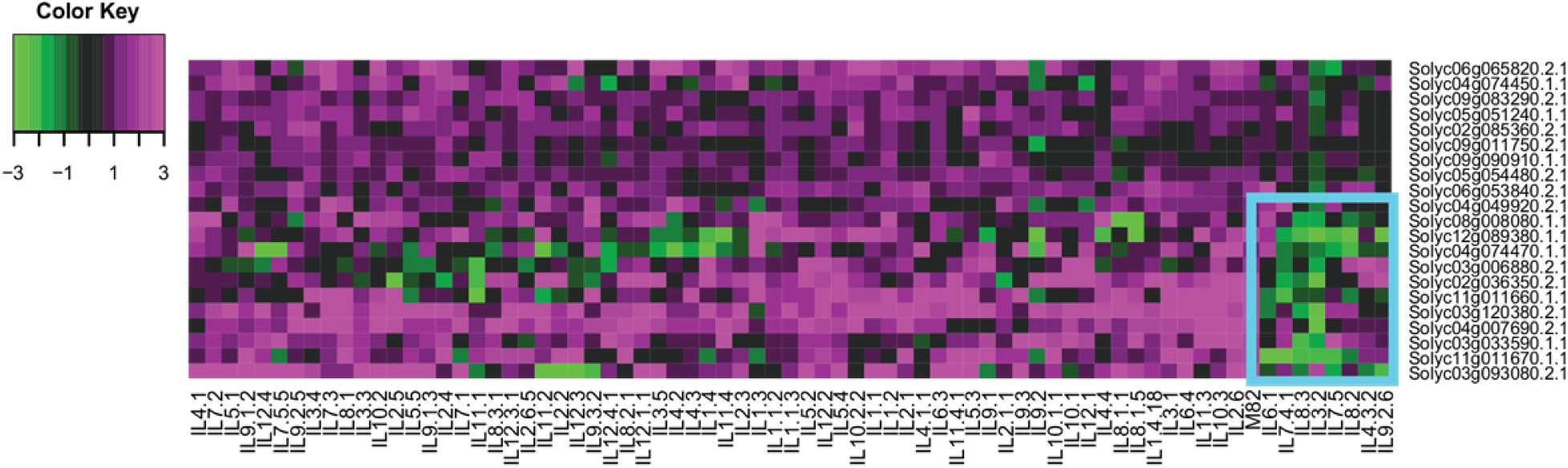
A heatmap displays the relative expression of a subset of shade responsive genes across the M82 × *S. pennellii* tomato introgression lines, clustered by Euclidean distance. Subset of auxin-associated genes (n = 21). Relative expression is calculated as the log2-transformed ratios of shade/sun expression values. Data were scaled within each IL prior to clustering. Green coloring represents genes where shade expression is less than that in sun (values < 0); magenta coloring represents genes > 0 where shade expression is greater than that in sun. Blue box indicates region of interest.

Returning to ILs of interest based on their phenotypes, we can now examine their gene expression patterns. Expression in IL9.3 and IL9.3.2 in response to shade differs in a number of auxin- and cell wall-related genes: IL9.3.2 has increased expression of auxin efflux carriers, *SAUR*s (for *SMALL AUXIN UPREGULATED*), *IAA* genes, and xyloglucanases and expansins, compared to IL9.3. These expression differences reflect the fact that IL9.3 is phenotypically much more shade tolerant than IL9.3.2. As described above, IL7.4.1 and IL3.2 do not respond strongly to shade in overall growth. This is reflected in reduced expression of shade-responsive auxin genes, shown in Fig. 7. IL11.4.1, which does not respond to shade in its petioles, has low expression of genes associated with growth in petioles; likewise, IL2.3 has minimal shade-responsive expression of petiole-related genes (see Fig. 8 and STable 8 below). Shade-responsive expression across the ILs is detailed in STable 3.

### Promoter enrichment analysis identifies auxin- and light-regulated motifs

Our findings in Fig. 7 and SFig. 8 led us to ask whether we find a particular set of motifs enriched in the putative promoters of the genes highlighted in each panel. We performed a promoter enrichment analysis on the subset of genes highlighted in SFig. 8. The binding sequence ofhomeotic transcription factor orthologs of the MADS-box AGAMOUS-LIKE family of proteins, AGAMOUS-LIKE 2/SEPELATA 1 (AGL2/SEP1) and AGL3/SEP4, were enriched in the promoters of the transcription factor cluster of genes (*p*-value < 0.05, STable 6, SFig. 8A). *AGL2* is typically known to be expressed in the floral meristem, whereas *AGL3* shows more broad expression in the aerial tissues (Ma et al., 1991; Gong et al., 2004), and both are involved in floral meristem determination. These differences in expression could relate to the changes in developmental rate induced by shade in this study. We also found marginal enrichment of two other promoter binding motifs: the homeodomain-leucine zipper II family member, *ATHB2* (*p-*value 0.1), and the Myb-related transcription factor *CIRCADIAN CLOCK ASSOCIATED1* (*CCA1*) (*p*-value <0.1). In Arabidopsis, *ATHB2* is regulated via *PHYTOCHROME INTERACTING FACTOR 4* (*PIF4*) and *PIF5* genes and is well known to activate genes implicated in the SAR (Hornitschek et al., 2009). *CCA1* is a central circadian clock oscillator, regulated by light (Wang et al., 1997; Alabadi et al., 2002).

The promoter motif enrichment search among the highlighted cell wall-associated genes (SFig. 8B) revealed enrichment for the binding site of *AGL1.* Although *AGL1* is commonly known for its involvement in flower development, it is moderately expressed in vegetative stem tissue as well (Ma et al., 1991; Gong et al., 2004). In agreement with the shade avoidance literature, the motif search revealed a marginally significant enrichment ofbinding sites of Auxin Response Factors (ARFs) in cell wall-associated genes (*p*-value < 0.1). Previous studies have linked increased auxin biosynthesis and signaling to the elongation of organs (Gray et al., 1998; Sessa et al., 2005; Tao et al., 2008; Sasidharan et al., 2014; Spartz et al., 2014).

Among the genes differentially expressed in the auxin cluster (Fig. 7, SFig. 8C), we found a significant enrichment (*p*-value < 0.01) for the I-box promoter motif (STable 6). In tomato, the I-box is the binding site for MYB1, a member of the R2R3 family of MYB-like transcription factors (Giuliano et al., 1988; Donald and Cashmore, 1990; Rose et al., 1999). This promoter sequence is conserved among the promoters of light-regulated genes in both Arabidopsis and tomato, including Rubisco subunits, and is homologous to the light regulated G-box binding domain (Giuliano et al., 1988). These findings point to the Conservation of targets of the shade avoidance response, which includes gene targets involved in light, auxin signaling and development.

### Genotypic and phenotypic correlations reveal genes differently regulated in petioles and internodes

We wanted to determine how well our study paralleled previously reported gene expression findings in tomato. A previous study by Cagnola and colleagues (Cagnola et al., 2012) examined the transcriptional response to shade in tomato leaves and stem tissue. We compared our list of shade-responsive tomato genes (Table 1; STable 2) to the shade-responsive gene expression found in the microarray experiment on *S. lycopersicum* leaves and stem segments grown in shade (Cagnola et al., 2012). Examining the genes common between our RNAseq analysis and the tomato microarray, nearly half of the genes identified as being shade responsive in the stem and leaf microarray study were also identified as being shade responsive in our data. The proportions of genes identified both by Cagnola and coworkers and by this study are shown in SFig. 7C (also see STable 7). Genetic shade responses of leaves and stems were examined separately by Cagnola and colleagues; our gene expression data is derived from tissue at the apical meristem including both leaf and stem primordia. However, we can examine leaf- and stem-specific shade responses by correlating gene expression with shade-responsive organ-specific phenotypes, such as internode elongation or petiole elongation.

Our phenotypic and PC analyses identified genotypes with organ-specific shade responses. These results prompted us to ask whether, like the work of Cagnola *et al.,* we could find genes that were significantly correlated with both internodes and petioles or specific to either of these organs. After performing correlations between gene expression and phenotype data, we mined the genes with a moderate to strong Spearman’s correlation coefficient, *rho* > |0.3|, that were identified to be correlated with either internode PC1 or petiole PC1 (Fig. 8A). We found that 157 genes were correlated with internode PC1 with a |*rho*| >0.3. Of these, 59 were similarly correlated with petiole PC1 (difference in *rho <* 0.15), whereas 98 showed a shigher correlation to internode PC1 (Fig. 8B). Similarly we observed that of the 54 genes correlated to petiole PC1, 31 showed a stronger correlation relationship with petiole PC1 than with internode PC1, 22 are similar, and 1 gene actually shows a higher correlation with internode PC1 (Fig. 8C). In other words, the majority of genes whose expression is associated with growth in internodes or petioles are associated only with growth in that specific organ. This result suggests the plant undergoes a modulation of organ-specific responses to shade, and reflects our findings of organ-specific SAR in the ILs. Further, the genes correlating with the internode and petiole PC1 traits reflect what is seen at a phenotypic level, in that there appears to be a stronger shade response in internodes than in petioles. For instance, we see higher correlation values for genes involving cell wall modification, hormone biosynthesis and hormone efflux in the internodes than we observe for the petioles (STable 8; see also SFig. 9 described below).

**Figure 8.**
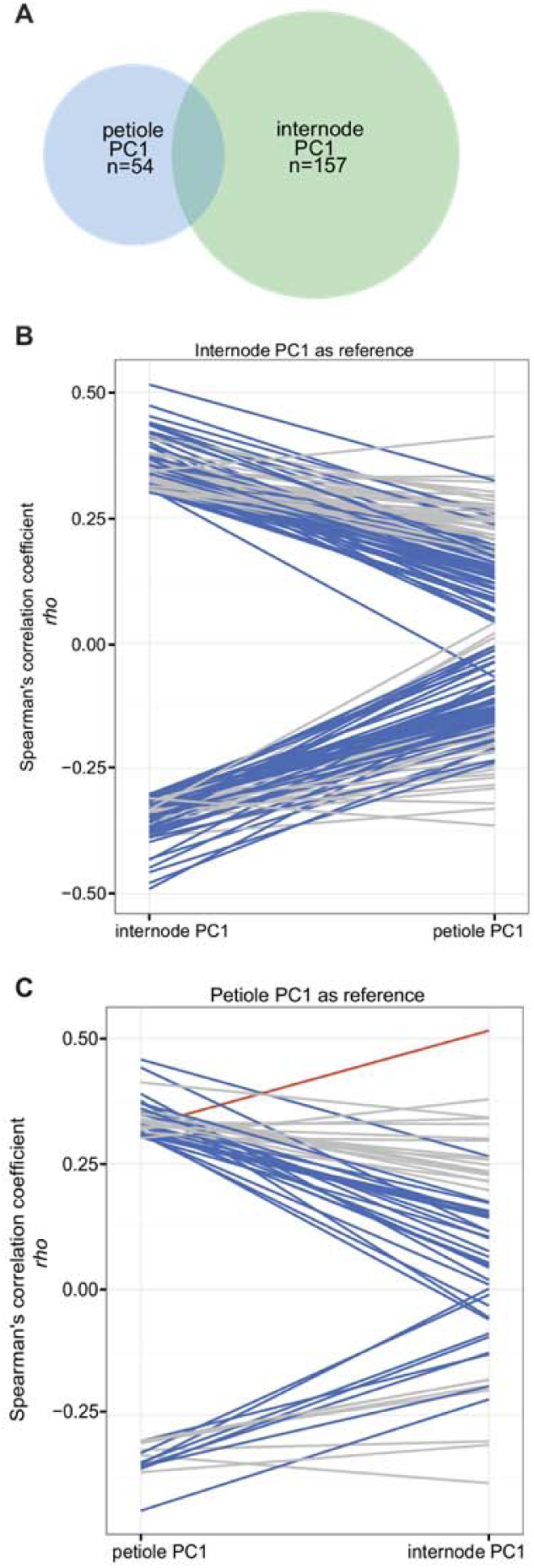
Organ specificity of gene and phenotype correlations. (A) Venn diagram showing the overlap between the genes correlated with internode and petiole PC1 data. (B) and (C) Reaction norm plots of correlation vatues between gene expression and phenotype measurements, with correlation greater than |0.3| with internode PC1 (B) or petiole PC1 (C). Blue lines indicate genes where the absolute value of rho is greater in the organ of interest; red lines indicate the reverse; gray lines indicate correlations that are similar in both organs (difference in *rho* between the organs < 0.15).

To expand our analysis of shade responsive gene expression and phenotype relationships, we performed phenotype-by-gene expression correlation analysis on all phenotypes and genes differentially expressed in shade. From these gene and phenotype correlations, we performed GO enrichment analysis. For each phenotype, we asked which functions might be over- or under-represented in the subset of correlated genes. These results are presented in SFig. 9 (see also STable 9). As expected, there is overlap between the correlated gene ontologies enriched in intPCl and total height. In contrast, several GO terms enriched in petPCl appear to be underrepresented in intPC2, and it may be of future interest to examine the relationship between leaf and stem in partitioning resources during a shade response. These oppositely regulated GO terms include several categories of glucosidases, suggesting a link to cell wall processes.

### Differential network analysis reveals few differences in network Connectivity

Given the large number of transcriptional changes induced by shade, we hypothesized that shade may alter the connectivity of genes within genetic networks. To test this idea, we performed a differential network analysis using the differential weighted co-expression network analysis (DWGCNA) function in the WGCNA R-package (Langfelder and Horvath, 2008). The DWGCNA method is based on the idea that expression of each gene is correlated with many other genes; the change in light treatment may result in a change in the networks of connected genes. After the connectivity for each gene (*k_i_*) was calculated in each network (sun and shade networks), the gene connectivity value was normalized by the average network connectivity for each respective network (*K*), followed by calculation of the difference between networks (Diff*K*).

The Diff*K* was calculated by subtracting the gene connectivity values for the shade network from the sun network. Fig. 9A presents a scatterplot with the *x*-axis displaying the differences in Connectivity (Diff*K*) between the sun and shade networks, and the *y*-axis representing differential gene expression shown as the paired *t*-statistic. This *t*-statistic was calculated for each gene by performing a *t*-test between the sun and shade expression across all the ILs. The nine sectors within the plot are defined by the boundaries applied to the *t*-statistic value (significance at *t >* |1.96|) and absolute values of the Diff*K* Connectivity score (>|0.2|). Genes found above *t* = 1.96 are more highly expressed in sun (sectors 1, 2, 3), and genes at *t* < 1.96 are more highly expressed in shade (sectors 5, 6, 7). Considering Diff*K*, genes with a connectivity greater than 0.2 display greater connectivity in sun (or, reduced Connectivity in shade; sectors 3, 4, 5), while genes with Diff*K* less than -0.2 have increased connectivity in shade (sectors 1, 7, 8). To test the significance of sector assignments, we permuted the “sun” and “shade” assignments across 1,000 data sets and compared the number of genes assigned to each sector. The number of genes in sectors 2, 3, 5 and 6 were determined to be significantly greater than expected (*p*-value < 0.01), suggesting a meaningful, non-random distribution of genes. The genes found in each of these sectors are presented in STable 10.

**Figure 9.**
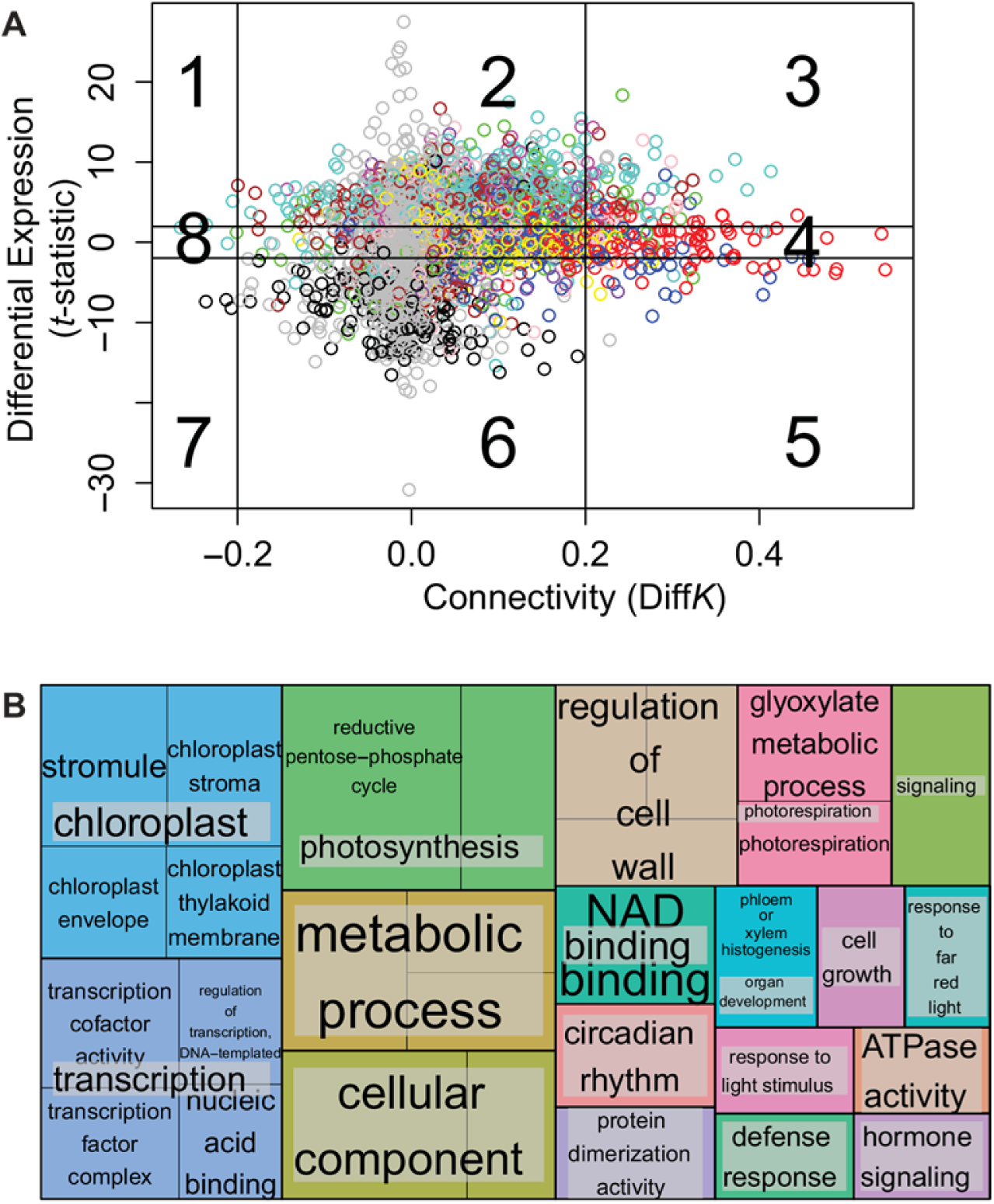
Differential gene network analysis (DWGCNA). (A) Differential network Connectivity between sun and shade gene networks. The x-axis plots the difference in Connectivity, with vertical lines indicating a difference of |0.2|. DiffK < 0 indicates greater Connectivity in shade. The y-axis show the t-statistic indicating expression differences between sun and shade; the horizontal lines delineate the t-statistic of |1,96|. T-statistic < 0 indicate genes more highly expressed in shade. These lines separate the plot into 9 sectors. Each circle represents a gene within the shade and sun networks, and is colored based on the gene ontology. (B) A REVI-GO tree map plot visualizes gene ontology lists of enriched categories from sector 3 (decreased shade Connectivity, decreased shade expression). The size of each box reflects both the p-value significance and number of terms fitting within a GO category.

Sectors 3 and 5 contain the genes that have the greatest connectivity differences between sun and shade networks. While connectivity between the networks does change with light treatment, shade treatment reduces the connectivity of only 7.5% of the genes in the network; this can be seen by comparing the density of genes found in sectors 3, 4, and 5 to the remaining sectors. Shade caused reduced connectivity of 7.3% of the genes with lower expression in shade (top row Fig. 9A), 9.8% of genes without differential expression (middle row), and 5.3% of the genes expressed more highly in shade (bottom row), indicating that shade is more likely to reduce connectivity among genes with decreased or no change in expression (p = 0.006, chi-squared test).

**Sector 3: Light and auxin signaling are responsible for changes in Connectivity between the sun and shade networks.** A gene ontology (GO) analysis of the genes in the co-expression network sector 3 confirmed that shade decreases connectivity among genes in several categories, including hormone signaling, light response, cell wall modification and photosynthesis (STables 10, 11). At first glance, it may be surprising to find that auxin signaling genes are enriched in this sector, with decreased expression and connectivity. It is well known that auxin-signaling is implicated in the shade avoidance response (Morelli and Ruberti, 2000; Carabelli et al., 2007; Tao et al., 2008; Ciolfi et al., 2013; de Wit et al., 2014). However, examination of the auxin-associated genes in SFig. 8C demonstrates that both up- and down-regulation of expression occurs in response to shade. For example, within the GO-enriched category ‘auxin signaling’, two Auxin Response Factor (ARF) transcription factors, *ARF8* and *ARF19,* are found in sector 3.

The enriched category ‘1,3-beta-D-glucan synthase complex’ contains genes including two *GLUCAN SYNTHASE-LIKE* genes, encoding callose synthase proteins. Callose synthases have diverse functions in response to abiotic and biotic stress, and increase callose deposition in cell walls within minutes of wounding or temperature change (Chen and Kim (2009); (Jacobs et al., 2003; Chen and Kim, 2009). While it is clear that callose synthases are important under stress scenarios, their role in shade avoidance is unknown.

Two additional sector 3 genes, Chlorophyll a/b binding protein and *ARGONAUTE1*, fall under the GO enriched category of ‘response to far-red light’. Chlorophyll a/b binding protein (CAB) is known to be regulated by phytochrome, with its expression under control of R and FR light in Arabidopsis seedlings (Karlinneumann et al., 1988; Dewdney et al., 1993; Casal and Yanovsky, 2005). In our gene expression study, a great proportion (approximately 70%) of the ILs show decreased expression of the CAB homolog in shade (SFig. 10A). *ARGONAUTE1* (*AGO1*) is a member of the essential catalytic components of the RNA-induced silencing complex, and plays an important role in plant development; *AGO1* is hypothesized to act as a link between light and auxin signaling (Kidner and Martienssen, 2004; Vaucheret et al., 2004; Sorin et al., 2005). *AGO1* mutants are hypersensitive to light, which may be a result of the upregulation of light signaling pathways. *AGO1* may also regulate auxin homeostasis in Arabidopsis by targeting the transcripts of *ARF17,* a repressor of auxin inducible genes (Sorin et al., 2005). If *AGO1* acts in tomato as it does in Arabidopsis, then *AGO1* may be acting as a positive regulator of the SAR by this suggested mechanism.

Another sector 3 GO enriched category is ‘response to light stimulus’. The GATA transcription factor and glyceraldehyde-3-phosphate dehydrogenase (GAPDH) are found in this category. Previous work has elegantly shown that components functioning in the Calvin cycle, which includes GAPDH, are light regulated (Dewdney et al., 1993; Tobin and Kehoe, 1994). Early work done in phytochrome gene regulation revealed that many genes regulated by phytochrome, including GAPDH, contain a GATA binding motif (Jeong and Shih, 2003).

**Sector 5: Chloroplast-related gene expression does not correlate with measured phenotypes.** Gene ontology enrichment analysis of the sector 5 genes indicated that this subset of genes are involved in chloroplast maintenance, protein transport and regulation of stromal and lumen reactions (STable 10). We examined the gene expression patterns of these genes in the ILs in shade, and found three trends in gene expression among the sector 5 genes (SFig. 10B). ILs clustered into approximately equal groups with strong up-regulation, strong down-regulation, or intermediate expression of sector 5 genes, though these clusters did not correspond directly to any measured shade-responsive phenotype. In this analysis, genotypes that are entirely contained within another IL tend to cluster together, for example IL9.3 and IL9.3.2 (Fig. 3D, SFig. 10B), as might be expected.

## Discussion

Knowledge of shade responses gamered from studies in *Arabidopsis thaliana* has served as a basis for understanding the shade avoidance response (SAR) in other species. However, these studies do not address agriculturally relevant types of shade avoidance, such as that in crop species with expanded vegetative internodes. Using a domesticated tomato and *S. pennellii* introgression line population, we were able to dissect the genetic basis of variation in the shade avoidance response in an important crop and its wild relative. As a relatively distant relative to the modern domesticated tomato, previous work has shown that *S. pennellii* gene expression has significantly diverged from that of domesticated tomato (Koenig et al., 2013). Such differences are evidence of the distinct evolutionary histories of the two species: the domesticated tomato was selected under favorable yet crowded conditions, whereas *S. pennellii* adapted to an arid and less populated environment. The IL population takes advantage of these species-specific differences, parsing the genome of the wild species into multiple individuals and separating inherent genetic interactions. Examining each IL and its associated *S. pennellii* genome region independently allows us to dissect the genetic basis for quantitative traits and complex genetic responses such as the SAR.

We found that variation in the SAR in this cross is genetically complex, involving multiple loci across the genome controlling distinct, yet quantifiable traits (Fig. 1, Fig. 3, SFig. 4). Corroborating previous results, we show that the tomato shade avoidance response is primarily modulated through internode elongation (internode PC1), as a greater number of ILs showed a significant increase in internode elongation than petiole elongation (petiole PC1). The differing degrees of phenotypic shade response observed in internodes and petioles suggest that the two organs have differing sensitivity to shade and that this may be reflected in their gene expression (Fig. 8). Further work quantifying expression of specific genes in internodes and petioles separately may validate this hypothesis.

A primary aim of this work was to examine shade-responsive gene expression in domesticated tomato using next-generation sequencing (Fig. 6, SFig. 6). RNA sequencing allowed for a more complete Identification of shade-responsive transcripts than could be provided in previous studies that used microarrays, such as in Cagnola *et al.* (Kidner and Martienssen, 2004; Vaucheret et al., 2004; Sorin et al., 2005). In combination with phenotypic analysis of growth in sun and shade conditions, our gene expression analysis allowed us to identify genes and regions of the genome critically involved in shade avoidance responses in tomato.

Comparison of our gene expression findings to previously reported work in Arabidopsis and tomato (Sessa et al., 2005; Tao et al., 2008; Cagnola et al., 2012) revealed both novel genes and homologs of previously characterized shade responsive genes (Table 1, STables 2 and 7). In Arabidopsis studies, the HD-ZIP II homeobox domain transcription factor *ATHB2* is a classic example of a shade-induced gene; it activates several shade avoidance response genes playing a role in elongation and growth (Carabelli et al., 2013). We found that each of the three tomato homologs of *ATHB2* (Solyc06g060830, Solyc08g078300 and Solyc08g007270) showed a significant up-regulation in shade (Table 1 and STable 2), in agreement with studies in Arabidopsis. Notably, the critical negative regulator of shade in Arabidopsis, *LONG HYPOCOTYL IN FAR RED 1* (*HFR1*), is not present in the tomato genome (Fankhauser and Chory, 2000; Sessa et al., 2005; Hornitschek et al., 2009). Our phenotypic measurements identified ILs with limited growth in shade, indicating there are a number of potential negative regulatory genes in the tomato genome; further work examining gene networks in sun and shade may reveal key shade regulators in tomato.

Though the comparison and contrast of our dataset to the aforementioned resources showed a number of genes commonly expressed in shade, a large number of the genes in our study were not identified in previous work (SFig. 7). We believe this difference to be due to differences in experimental design. Environmental conditions, developmental stage, and sampled tissue can have a great impact on gene expression results. In our study, we used a R:FR ratio of 0.5 for our simulated shade condition, whereas Cagnola and colleagues (2012) performed their study using a low R:FR ratio of 0.05. The work in tomato by Cagnola and colleagues also looked at shade treatments of 1 hour and 4 days, contrasted with our shade treatment of 28 hours. Furthermore, the size of our experiment resulted in much greater power to detect expression differences, as we had 5 replicates of more than 80 samples in both sun and shade treatments, while the Arabidopsis studies used only 3 replicates in each light treatment (Sessa et al., 2005; Tao et al., 2008). Altogether, these distinctions explain the significance of differences in the shade-responsive genes identified between studies.

Clustering the ILs based on their shade-responsive gene expression patterns led us to focus on three subsets of genes, namely transcription factors, auxin-related genes, and cell wall-related genes (SFig. 8). A small group of transcription factors is upregulated in the ILs during shade; this includes each of the three *ATHB2* homologs (SFig. 8A). The transcription factor *ESE3* has been identified as shade responsive in Arabidopsis, but the other seven genes, including MADs box, Myb, and bHLH transcription factors, have not previously been shown to play a role in shade-responsive growth. A group of ten ILs clustered together based on their altered cell wall-related gene expression (SFig. 8B); the differentially expressed cell wall genes included a number ofhydrolases and proteins with polysaccharide transferase activity. While cell wall-associated genes were correlated with shade-responsive growth (STable 9), we were unable to identify a consistent phenotypic trait altered in ILs with distinct cell-wall gene expression patterns; future experiments may target cellular-level rather than organismal-level traits, such as cell wall composition, in order to connect expression of these genes with a specific growth phenotype. Considering the auxin-related genes, a group of eight ILs cluster based on differential expression of 12 genes (Fig. 7, SFig. 8C). Among typical auxin-responsive genes, this list also includes expansins, *XTH* genes, and *SAUR* genes, which have all been shown to be involved in auxin-mediated cell wall expansion (Table 3). Importantly, the eight ILs clustering based on their auxin gene expression also cluster based on their shade-responsive intemode elongation (intemode PC1) phenotype (Fig. 7, SFig. 4C). Together these eight ILs contain six non-overlapping regions, indicating that these auxin-responsive genes are a common downstream target among independent QTL promoting reduced shade avoidance response.

Based on the clusters of shade-responsive transcription factor, auxin, and cell wall genes, we also performed an analysis for enrichment of promoter motifs. In accordance with our previous findings, promoters of the transcription factors are enriched for ATHB2 binding sites, supporting light-regulated modulation of expression. Auxin genes differentially expressed in shade show enrichment of light-signaling promoter elements, while the promoters of cell wall genes are enriched for auxin-related binding sites.

The results described above allowed us to determine whether shade alters the Connectivity of the gene network. Using DWGCNA, a differential network analysis method (Fuller et al., 2007; Langfelder and Horvath, 2008), we found that the sun and shade gene correlation networks are most contrasted by their differences in gene expression, seen in sectors 2 and 6 in Fig. 9A. However, shade does alter the network connectivity of a number of genes, found primarily in sectors 3 and 5. Our analysis of sector 3 showed that within this sector are included genes involved in light and hormone signaling and photosynthesis, though we could not relate that pattern of sector 3 gene expression to ILs that were particularly shade sensitive or shade tolerant. Sector 5, in which connectivity of genes decreases in shade, includes a number of chloroplast-related genes; combined with the sector 3 information, this data suggests that after 28 hours of shade growth, regulation of photosynthetic gene expression is altered. Examination of gene expression after longer periods of shade exposure may reveal reduced photosynthetic signaling, as previous physiological studies have suggested (Boardman, 1977).

All told, this large-scale study of shade from both phenotypic and gene expression standpoints has provided us with an unprecedented data set for identifying key genes involved in general and organ-specific shade responses in tomato. The environment has a profound influence on plant development, modulating growth for optimal capture of resources, sometimes to the detriment of plant productivity. Especially relevant for photosynthetic organisms, neighbor-shading can modify plant growth to optimize light acquisition thus impacting agricultural yield. In this paper, we have only begun the analysis of the relationships between natural variation in shade-responsive phenotypes and gene expression. In combination with the previously studied Arabidopsis and tomato work, this work may enable development of more shade-tolerant varieties of tomato throughjudicious use of wild and domesticated tomato germplasm as well as genomic resources.

## Methods

### Plant growth conditions

**Germination.** The introgression line population was obtained from the UC Davis/C.M. Rick Tomato Genetics Resource Center and maintained by the Department of Plant Sciences, University of California, Davis, CA. Seeds were scarified with 50% household bleach for 5 minutes and rinsed five times with sterile MilliQ water prior to sowing onto moist paper towels housed in phytatrays (Sigma-Aldrich, part no. P5929). The seeds were allowed to germinate in the dark at room temperature for three days. At the end of this period, the germinated seedlings were transferred to ‘sun’ conditions, lights described below, for four days prior to transplantation. At seven days after seed imbibition (dai), the fully germinated and emerged seedlings were transplanted into 620mL capacity pots with commercial Sunshine Mix No. 1 (Sun Gro Horticulture). The seedlings were watered with Grow More Inc. 4-18-38 hydroponic solution. Upon transplantation, the seedlings were placed in their respective light treatment. The experiment was concluded at35 dai.

**Chamber and light conditions.** The 76 introgression line population was grown in chamber conditions under 16h light and 8h dark cycles at 22°C. To achieve the simulated sun and shade conditions, we used cool white fluorescent and infra-red (IR) bulbs (F48T12 VHO far red bulbs, Interlectric Corp., Warren PA). We achieved approximately 110 micromoles of photosynthetically active radiation in both sun and shade treatments. For the simulated shade conditions, fluorescent and IR bulbs produced a low R:FR ratio of 0.5; in simulated sun conditions, we placed filters over the IR bulbs to reach a high R:FR of 1.5.

### Growth measurements and analysis

**Experimental design.** The IL population, including *Solanum pennellii* and *Solanum lycopersicum* cv. M82, was represented once per replicate, achieving a total of 11 individuals of each genotype per treatment through eleven replicates. We grew three replicates at a time. All individuals in each replicate were randomized and mirrored in each light treatment. Our analysis of the IL growth data included a linear model accounting for position, shelf and temporal replication as random effects.

**Measurement of internode and petiole traits.** Internode lengths between adjacent leaf axils were measured for plants grown in sun and shade using Vernier digital calipers. Epicotyl and internodes 1 to 3 were measured at 28 dai (4 weeks) while the epicotyl and all visible internodes were measured at35 dai (5 weeks). Internodes were counted from the bottom up, with the epicotyl representing the region of the stem between the cotyledons and first leaf node, and internode 1 being the region between leaf 1 and leaf 2. Total plant height, or length of the primary stem, was measured at 35 dai, from the cotyledons to the tip of the SAM. Petiole length measurements on 35 dai were performed on images of the lowermost four leaves of each plant. The leaves were cut at the stem and subsequently imaged (Chitwood et al., 2014). The petioles were measured using the ImageJ Software (Abramoff MD, 2004), starting at the cut end of the petiole to the first proximal leaflet.

**Principal component analysis, linear modeling of traits, and visualization of phenotypic data.** Intemode and petiole measurements, as well as the growth measurements calculated from week 4 and week 5 data, were each subjected to principal component analysis with the statistical program R (R: A language and environment for statistical computing., 2015). Data were scaled and centered, and the principal components (PCs) were calculated using the `prcomp()` function in the basic `stats` package in R. Loadings such as those in Fig. 2 were generated from the rotation content in the PC analysis.

Mean values and Standard errors for all traits, including untransformed traits such as internode number as well as PCs, were calculated using the `lmer()` function in the `lme4` (linear mixed-effects models using Eigen and S4) package in R (Pinheiro J, 2014). P-values were calculated to indicate statistical difference from the control M82. Standard errors of the log_2_-transformed ratio of shade/sun values were generated using `lsmeans()` in the `lmerTest` package (Bates D, 2014).

Plots of trait and PC values such as those in Fig. 3 and SFig. 3 were generated using the `ggplot2` package in R (Wilkham, 2009). Mean values and Standard errors were incorporated using the `geom_point()` and `geom_linerange()` functions. Piecharts such as those in SFig. 6 were generated using the ggplot functions `geom_bar(stat = "identity")` and `coord_polar()`.

**Bin mapping via regularized regression.** For each trait analyzed a linear mixed effect model was fit using the `lmer()` function from the `lme4` package (Bates D, 2014), with replicate and shelf as random effects and IL, treatment, and their interaction as fixed effects. The predicted random effects were extracted from the model fit and used to adjust the raw observations. For bin mapping we then used the `cv.glmnet()` function from the `glmnet` package (Wilkham, 2009), with bin genotypes, treatment, and the genotype by treatment effect as predictors. The mixing parameter alpha was set at 0.6 and 100-fold cross validation was used to choose an appropriate value for the penalization factor lambda.

### Transcriptome sequencing and analysis

**Library preparation.** We sequenced the transcriptomes of all the ILs, including the parents *S. lycopersicum* cv. M82 and *S. pennellii* for after treatment with sun and shade conditions. The plants were germinated and transplanted as described above. Plants were grown for 18 days in sun; all plants were then transferred to a second chamber with identical growth conditions for a 28-hour shade or sun treatment prior to sampling of tissues. Tissues above the cotyledons were sampled, including the epicotyl, apical meristem, and leaf primordia. Expanding leaves greater than 0.5cm in length were excluded. The RNA-seq libraries were prepared and sequenced as described by Kumar and colleagues(Kumar et al., 2012). The next-generation sequencing reads were quality-filtered and trimmed using the FastX-Toolkit (Hannon, 2009) and customized perl scripts. Adapter sequences were removed and reads were sorted by their sample-specific barcodes. Reads were next mapped to tomato reference sequences as described in Chitwood et al. (2013) (Kumar et al., 2012). Based on the read mapping, gene-specific counts were calculated for each sample and used for differential expression analysis.

**Differential gene expression analysis.** To perform differential gene expression analysis, we used the statistical analysis program R and the package `‘edgeR’` from Bioconductor (Robinson et al., 2010). EdgeR uses empirical Bayes estimation and an exact test based on the negative binomial distribution for use with next-generation sequencing read count data. This allows for the differential analysis of gene expression data from RNA-seq experiments that include biological replication (Robinson et al., 2010). We began with a list > 30,000 tomato genes from the Heinz 1706 genome assembly (Consortium, 2012). The genes were filtered to include only those genes which had a value of 5 counts per million (cpm) in at least 5 samples. Normalization factors used to scale each library were generated using the TMM method in `calcNormFactors()`, and dispersion values were calculated for each gene using edgeR’s function `estimateGLMTagwiseDisp()`. To generate the list of approximately 8000 genes differentially expressed in shade, we used a generalized linear model (edgeR functions `glmFit()` and `glmLRT())` to identify differential expression of genes between plants grown in sun or shade conditions, ignorant of the genotype of any plant. Genes differentially expressed in a genotype-specific manner, as well as genes with genotype-by-treatment interaction effects, were also identified using the generalized linear model in edgeR.

**Calculation of enrichment of gene ontology classes.** With the `goseq` and `GO.db` packages for R, we employed the tomato gene ontology categories (Koenig et al., 2013) to calculate enrichment of ontologies in our gene lists of interest using the `nullp()` and `goseq()` functions (Young etal., 2010; M, 2014).

**Network building.** Gene expression data were first processed using the `VOOM` R-package, allowing for the transformation of count expression data to log2-counts per million, and therefore permit estimation of the mean-variance relationship to compute appropriate observational-level weights (Law et al., 2014). Upon transformation, the shade avoidance response gene expression was calculated by subtracting log2(shade) from log2(sun) VOOM-transformed data. The gene expression datasets were reduced by finding genes with common high Connectivity in both datasets using the `softConnectivity()` function in the `WGCNA` (Weighted Gene Co-expression Analysis) R package (Langfelder and Horvath, 2008). This procedure yielded approximately 3,100 genes that were then used to move forward with the differential weighted gene co-expression network. This method of filtering allowed us to raise the adjacency to a common power of 14 to build the co-expression network. We then followed the DWGCNA procedure as described in (Fuller et al., 2007). In short, Diff*K* and *t*-statistic values were calculated to identify genes with large and significant differences in connectivity and expression between sun and shade networks. Plotting Diff*K* and the *t*-statistic, we assigned significance levels (*p*-values) to each sector of the scatterplot by performing 1000 random permutations within each sector. The permutation test contrasts the networks built by randomly partitioning the gene expression provided by the individual genotype samples in both aforementioned conditions. Based on the permutations, sectors 2, 3, 5 and 6 were found to be significant with a *p*-value < 0.01. The significant sectors were subjected to GO enrichment analysis as described above.

**Motif enrichment.** To perform the promoter enrichment search, we used 1000 bp upstream of the start codon of each subset of genes of interest as highlighted in Fig. 7. We then searched for the total number of incidences of any motif within the given target sequence and performed a Fisher’s exact test to calculate significance.

**Genotype and phenotype correlation analyses.** For each IL, mean values for gene expression and trait measurements were explored for meaningful associations. Relationships between genotype and traits such as those seen in Figs. 4 and 7 were calculated using the `dist()` function to determine Euclidean distance. Relationships between gene expression and phenotype such as those used in SFig. 9 were calculated using `cor.test(method = "Spearman")`. Data values were scaled but not centered in order to maintain any relationship around zero, followed by the `hclust()` function in the basic R `stats` package. Plots were visualized using the `gplots` package `heatmap.2()` function (Warnes GR, 2009).

## Acknowledgements

We thank Dr. José A. Aguilar-Martínez, Enrique Ostria Gallardom, Kristina Zumstein, Maxwell Mumbach, Sony Daggupati, and Ashlee Gamarra for their essential work in planting and collecting raw data during the course of this experiment. LGC was funded through a National Institute of General Medical Sciences Training Grant (T32-GM007377).

## Supporting Data

### Supplemental Figures

**Supplemental Figure 1. Growth of wild and domesticated tomato species in sun and shade.** Values of (shade/sun) growth are plotted for total plant height and lengths of the hypocotyl, internode 1, and intemode 2. Cultivated and wild species display differences in shade response both within and between categories of tomato.

**Supplemental Figure 2. Internode and petiole measurements are highly correlated in the M82 × *S. pennellii* tomato introgression population.** Scatterplots of internode and petiole measurements reveal many positive correlations.

**Supplemental Figure 3. Principal component analysis of shade-related traits.** (A) For each trait examined, PC1 explains > 50% of the variation seen in the data, while PC2 explains approximately one-quarter of the variation. (B) and (C) Loadings of PC1 (B) and PC2 (C) across each of the organs measured at week 4. (D) and (E) Loadings of PC1 (D) and PC2 (E) across each of the organs for which a one-week growth rate was calculated.

**Supplemental Figure 4. Means and Standard errors of the relative shade response for each trait in the M82 × *S. pennellii* tomato introgression lines (log_2_-transformed ratios of shade/sun).** (A) internode number, (B) total height, (C) intemode PC1, (D) internode PC2, (E) early internode PC1, (F) early intemode PC2, (G) growth rate PC1, (H) growth rate PC2. *P-*values represent the significance compared to M82. (I) Scatterplots of the relative shade response of internode PC2, petiole PC2, and intemode number reveal strong positive correlation between the traits (Pearson’s *rho >* 0.5).

**Supplemental Figure 5. Sun- and shade-responsive traits are mapped to bins within the introgression lines.** We determined the contribution of each bin to a given phenotype using elastic net regularized regression. A value equal to 0 indicates the bin does not contribute to the phenotype. The contributions of each bin to each phenotype are shown: (A) intemode PC1 in sun, (B) petiole PC1 in sun, (C) intemode PC2 in sun, (D) internode PC2 in shade, (E) petiole PC2 in sun, and (F) petiole PC2 in shade. Circles represent no effect on phenotype and triangles represent a non-zero effect of the bin.

**Supplemental Figure 6. A heatmap displays the relative expression of all the shade responsive genes (n = 8352) across the M82 × S. pennellii tomato introgression lines, clustered by Euclidean distance.** Relative expression is calculated as the log2-transformed ratios of shade/sun expression values. Data were scaled within each IL prior to clustering. Green coloring represents genes where shade expression is less than that in sun (values < 0); magenta coloring represents relative expression > 0 where shade expression is greater than that in sun.

**Supplemental Figure 7. Novel and conserved genes are differentially expressed in response to shade in tomato.** (A) Arabidopsis homologs of shade-responsive tomato genes (*n* = 8352) include genes that have been published as part of specific processes (62%) as well as genes that have not been identified in previous studies (38%). Genes previously published as part of light or shade response make up 39% and 8.5% of shade-responsive tomato genes, respectively. (B) The subset of shade-responsive tomato genes homologous to Arabidopsis genes present on the ATH microarray (*n* = 4574) were compared to previous microarray studies of Arabidopsis shade response (Tao *et al* 2008; Sessa *et al* 2005). Together, the Arabidopsis studies identified just over 16% of the genes identified by our tomato study. (C) The shade responsive tomato genes we identified were subset to include only those present in the tomato genome microarray (*n* = 4193) used in the study by Cagnola *et al* (2012). Genes from Cagnola’s data were grouped as being equally shade-responsive in both leaves and stems, or more shade-responsive in either stem or leaf tissue; in total, these categories comprised 44% of the genes identified in our tomato study.

**Supplemental Figure 8. Heatmaps display the relative expression of subsets of shade responsive genes across the M82 × *S. pennellii* tomato introgression lines, clustered by Euclidean distance.** (A) Subset of genes classified as transcription factors in Arabidopsis (*n* = 103). (B) Subset of cell-wall associated genes (*n* = 125). (C) Subset of auxin-associated genes (*n* = 125). Relative expression is calculated as the log_2_-transformed ratios of shade/sun expression values. Data were scaled within each IL prior to clustering. Green coloring represents genes where shade expression is less than that in sun (values < 0); magenta coloring represents genes > 0 where shade expression is greater than that in sun. Blue boxes indicate regions of interest.

**Supplemental Figure 9. Phenotype-specific gene ontology categories.** A heatmap displays the significant enrichment of gene ontology (GO) categories correlated with each phenotype. GO categories over-represented in the data are indicated by magenta coloring, while under-representation is green. Color scaling is based on the -log_2_-transformed *p*-value associated with the GO enrichment test. Supplemental Table 9 presents this data numerically.

**Supplemental Figure 10. A heatmap displays the relative expression of genes in DWGCNA sectors 2, 3, 5 and 6 across the M82 × *S. pennellii* tomato introgression lines, clustered by Euclidean distance.** (A) Subset of genes in sector 3, for which Connectivity increases in shade. (B) Subset of genes in sector 5, for which Connectivity decreases in shade. (C) Subset of genes in sector 2, for which expression increases in shade. (D) Subset of genes in sector 6, for which expression decreases in shade. Relative expression is calculated as the log_2_-transformed ratios of shade/sun expression values. Data were scaled within each IL prior to clustering. Green coloring represents genes where shade expression is less than that in sun (values < 0); magenta coloring represents relative expression > 0 where shade expression is greater than that in sun.

### Supplemental Tables

**Supplemental Table S1.** Chromosome 2 genes within IL2.3 located in bins d-2E, d-2F, and d-2I. *ITAG, S. lycopersicum* gene identifier. *Chromosome,* location of each gene within the tomato genome. *Bin,* location of each gene within the bins of the M82 × *S. pennellii* tomato introgression population. *AGI,* Arabidopsis gene identifier. *Gene description,* published description of gene or homologs.

**Supplemental Table S2.** Genes identified as significantly responsive to shade in tomato. *ITAG, S. lycopersicum* gene identifier. *logFC,* log_2_ fold change of expression in shade compared to values in sun. *logCPM,* log_2_ counts per million reads across all samples. *LR,* likelihood ratio; large values indicate a greater likelihood of differential expression. *PValue,* significance of differential expression. *FDR,* false discovery rate calculated by adjusting the p-value using FDR < 0.01. *AGI,* Arabidopsis gene identifier. *Gene name,* Arabidopsis gene name. *Gene description,* published description of gene or homologs.

**Supplemental Table S3.** Genes identified as differentially expressed in the IL population in sun and shade. *ITAG*, S. lycopersicum gene identifier. *IL*, introgression line coefficient of expression in sun compared to M82 in sun. *SLY_shadeResp*, M82 coefficient of expression in shade compared to M82 in sun. *IL_shadeResp,* introgression line interaction coefficient of expression in shade. *logFC,* log2 fold change of expression in shade compared to values in sun. *logCPM,* log2 counts per million reads across all samples. *LR,* likelihood ratio; large values indicate a greater likelihood of differential expression. *PValue,* significance of differential expression. *FDR,* false discovery rate calculated by adjusting the p-value using FDR < 0.01. *AGI,* Arabidopsis gene identifier. *symbol,* Arabidopsis gene name. *gene_name,* published description of gene or homologs.

**Supplemental Table S4.** *Arabidopsis thaliana* genes used in analysis of tomato shade-responsive gene expression. Column names indicate the publication from which the gene list was obtained.

**Supplemental Table S5.** Gene ontology categories enriched in those shade-responsive tomato genes which were also found in the studies done by Sessa *et al.* (2005) and Tao *et al.* (2008). *GO hierarchy,* ontology category: MF, molecular function; CC, cellular component; BP, biological process.

**Supplemental Table S6.** Motifs identified as being over-represented in the promoters of shade-responsive genes (Fig. 7). *Percent, universal,* the expected frequency of observing a motif across the genome. *Percent, IL cluster,* the actual frequency of observing the motif in the IL gene clusters shown in Fig. 7. *p-value,* Fisher’s exact test significance value indicating the difference between universal and IL cluster percent values.

**Supplemental Table S7.** Shade responsive tomato genes identified in this study as well as in Cagnola *et al.* (2012). *ITAG, S. lycopersicum* gene identifier. *DE,* experiment in which the gene was differentially expressed: shade, shade-responsive in both leaves and stems; leaf, shade-responsive in leaves; stem, shade-responsive in stems. *AGI,* Arabidopsis gene identifier.

**Supplemental Table S8.** Organ-specific shade responsive genes, based on correlation of genes with Internode PC1 and Petiole PC1. *ITAG, S. lycopersicum* gene identifier. *rho_petiolePC1* and *rho_internodePC1,* Spearman’s correlation *rho* value between each phenotype and genotype. *Primary phenotype,* Phenotype of initial observation of *rho >* |0.3|. *AGI,* Arabidopsis gene identifier. *Gene name,* Arabidopsis gene name. *Gene description,* published description of gene or homologs.

**Supplemental Table S9.** Gene ontology categories enriched in genes significantly correlated with shade-responsive phenotypes. The column *Differential expression of GO category* indicates whether the GO category was up-regulated and enriched, or down-regulated and depleted. *log2_updn* incorporates the log_2_-transformation of the p-value with the positive or negative state indicated (*Differential expression of GO category*), utilized in SFig. 9.

**Supplemental Table S10.** Genes clustering into sectors 2, 3, 5, and 6 based on DWGCNA clustering. *ITAG, S. lycopersicum* gene identifier. *DWGCNA Sector,* sector in which each gene is found: sector 3 genes show an increase in Connectivity in shade; sector 5 genes show a decrease in Connectivity in shade; sector 2 genes have increased expression in shade; sector 6 genes have decreased expression in shade.

**Supplemental Table S11.** Gene ontology categories over- or under-represented in DWGCNA sectors 2, 3, 5, and 6. *DWGCNA Sector,* sector with which each gene ontology term is associated. *GO hierarchy,* ontology category: MF, molecular function; CC, cellular component; BP, biological process. *Adjusted p-value* represents the *p-value* adjusted using FDR < 0.01. *Differential expression of GO category* indicates whether the GO category was up-regulated and enriched (“up”), or down-regulated and depleted (“dn”). *Number of Differentially Expressed Genes in GO Category* and *Number of Genes in GO Category* present the observed and total number of genes within a GO category in each sector.

**Supplemental Table S12.** Expression of Auxin-related genes in shade. ILs and genes were subset based on Figure 7C. *Auxin Gene,* gene identifier. *Gene Description,* published description of gene or homologs. *Gene Location,* location of each gene within the *S. pennellii* ILs. *LogFC Expression,* log_2_ fold change of expression in shade compared to values in sun within each IL.

## Footnotes

This work was funded through a National Science Foundation grant (IOS-0820854) to Neelima Sinha, Julin Maloof, and Jie Peng.

